# Dynamical flexible inference of nonlinear latent structures in neural population activity

**DOI:** 10.1101/2023.03.13.532479

**Authors:** Hamidreza Abbaspourazad, Eray Erturk, Bijan Pesaran, Maryam M. Shanechi

## Abstract

Inferring complex spatiotemporal dynamics in neural population activity is critical for investigating neural mechanisms and developing neurotechnology. These activity patterns are noisy observations of lower-dimensional latent factors and their nonlinear dynamical structure. A major unaddressed challenge is to model this nonlinear structure, but in a manner that allows for flexible inference, whether causally, non-causally, or in the presence of missing neural observations. We address this challenge by developing DFINE, a new neural network that separates the model into dynamic and manifold latent factors, such that the dynamics can be modeled in tractable form. We show that DFINE achieves flexible nonlinear inference across diverse behaviors and brain regions. Further, despite enabling flexible inference unlike prior neural network models of population activity, DFINE also better predicts the behavior and neural activity, and better captures the latent neural manifold structure. DFINE can both enhance future neurotechnology and facilitate investigations across diverse domains of neuroscience.

## Introduction

Neural population activity exhibits rich spatiotemporal dynamical patterns that underlie our behaviors and functions^1–16^. Developing precise data-driven models of these complex dynamical patterns is critical both to study the neural basis of behavior and to develop advanced neurotechnology for decoding and modulation of brain states^17, 18^. Given the spatiotemporal correlations in population activity, how this activity evolves in time can be modeled more efficiently in terms of lower-dimensional latent factors.

These factors can lead to new scientific discovery by revealing new low-dimensional structures in coordinated population activity, which are not directly evident from the high-dimensional activity itself or from single-unit activities^1–23^. These latent factors can also decode behavior to enable enhanced neurotechnology and brain-machine interfaces (BMIs)^17, 23^. As such, a critical objective is to develop latent factor models that not only are accurate in characterizing neural population activity with potential nonlinearities, but also enable the flexible inference of latent factors so that these factors can be seamlessly extracted in basic science and neurotechnology applications. Despite much progress, this objective of simultaneously enabling both accurate nonlinear modeling and flexible inference has remained elusive.

First, to be accurate, such a model should (i) capture potential nonlinearities in neural population activity and (ii) have a data-efficient architecture for generalizable data-driven training. Second, to enable flexible inference, such a model should (iii) be capable of both causal/real-time and non-causal inference simultaneously and (iv) allow for inference in the presence of missing neural measurements. If achieved, flexible inference will allow the same trained model not only to operate causally and recursively in real-time to enable neurotechnology, but also to leverage the entire length of data non- causally for more accurate inference in scientific investigations. Such inference will also make the model robust to missing or noisy measurements. Current dynamical models of neural population activity do not satisfy all the above four properties simultaneously.

Many of these models are linear or generalized linear, often in the form of linear dynamical models (LDMs)^1, 2, 11, 12, 21, 24–27^, and are used to infer low-dimensional latent factors^1, 11, 12, 21, 24–27^ or build BMIs^2^ (see also **Discussion**). However, while LDMs are data-efficient to train and allow for flexible inference with Kalman filtering^28^, they cannot capture the potential nonlinearities in the neural population activity to describe it more accurately. Beyond linear models, recent studies have leveraged the richness of deep learning to develop generative dynamical models of neural population activity^4, 13, 29–32^. However, while these models can capture nonlinearity, they do not meet all the flexible inference properties outlined above. Because the inference for these models is not solvable analytically unlike LDMs, they need to empirically train an inference or recognition network simultaneously with their generative network, usually requiring the entire length of data over a trial. Thus, their inference depends on how the specific inference network is structured and is not flexible to satisfy all the above properties. Indeed, in prior generative models, including sequential autoencoders^4^ (SAEs) or LDMs with nonlinear embeddings^30^, inference is non-causal, and real-time and/or recursive inference is not directly addressed^4, 13, 29, 30, 32^.

Further, in these models, inference in the presence of missing observations is not directly addressed^4, 13, 30–32^ and using zeros instead of missing observations^33, 34^ can yield sub-optimal performance by changing the inherent values of missing observations during inference^35, 36^ (see **Results** and **Discussion**). Similarly to these generative models, predictive dynamical models of neural activity that use forward recurrent neural networks^37–39^ (RNNs) also do not enable the flexible inference properties above. While these models can perform causal inference, they do not allow for non-causal inference to leverage all data, and they do not directly address inference with missing observations similar to above generative models.

Here, we develop a neural network model that encompasses both flexible inference and accurate nonlinear description of neural population activity. To achieve this, we build a network architecture consisting of two sets of latent factors rather than one. One set termed dynamic factors captures how neural population activity evolves over a low-dimensional nonlinear manifold, and the other set termed manifold factors characterizes how this manifold is embedded in the high-dimensional neural activity space. By separating these two sets of factors, we can capture the nonlinearity in the manifold factors while keeping the dynamics on the manifold linear, thus enabling flexible inference by exploiting the Kalman filter on the nonlinear manifold (see **Methods** and **Discussion**). We term this method Dynamical Flexible Inference for Nonlinear Embeddings (DFINE).

We validated DFINE in nonlinear simulations and then compared it to benchmark linear LDM and nonlinear SAE methods across diverse behavioral tasks, brain regions and neural signal types. We found that DFINE not only enabled flexible inference capabilities in nonlinear modeling, but also performed significantly more accurately than both linear and nonlinear benchmarks. First, given its flexible inference, DFINE robustly compensated for missing observations and seamlessly performed both causal and non-causal inference of latent factors. Second, compared to the linear and nonlinear benchmarks, DFINE significantly improved the accuracy in predicting neural activity and behavior, and in recovering the low-dimensional nonlinear neural manifold in single trials. Finally, we extended DFINE to supervise the model training with behavior data, such that the extracted latent factors from neural activity are more predictive of behavior. DFINE enables the new capability of flexible inference in nonlinear neural network modeling and enhances neural description for diverse neurotechnology and neuroscience applications.

## Results

### Summary of methods

We develop DFINE as a novel nonlinear neural network model of neural population activity with the new capability to perform flexible inference. We also devise the associated learning and inference methods. To model neural population activity, we define two sets of latent factors: the dynamic latent factors which characterize the temporal dynamics on a nonlinear manifold, and the manifold latent factors which describe this low-dimensional manifold that is embedded in the high-dimensional neural population activity space (**Fig. 1a**). This separation allows the model to capture nonlinearity with the link between the manifold factors and neural population activity, while keeping the dynamics on the manifold linear (**Discussion**). As such, these two separate sets of latent factors together enable all the above flexible inference properties by allowing for Kalman filtering on the manifold while also capturing nonlinearity. This flexible inference includes the ability to perform both causal (filtering) and non-causal (smoothing) inference, and to perform inference in the presence of missing observations.

**Figure 1.**
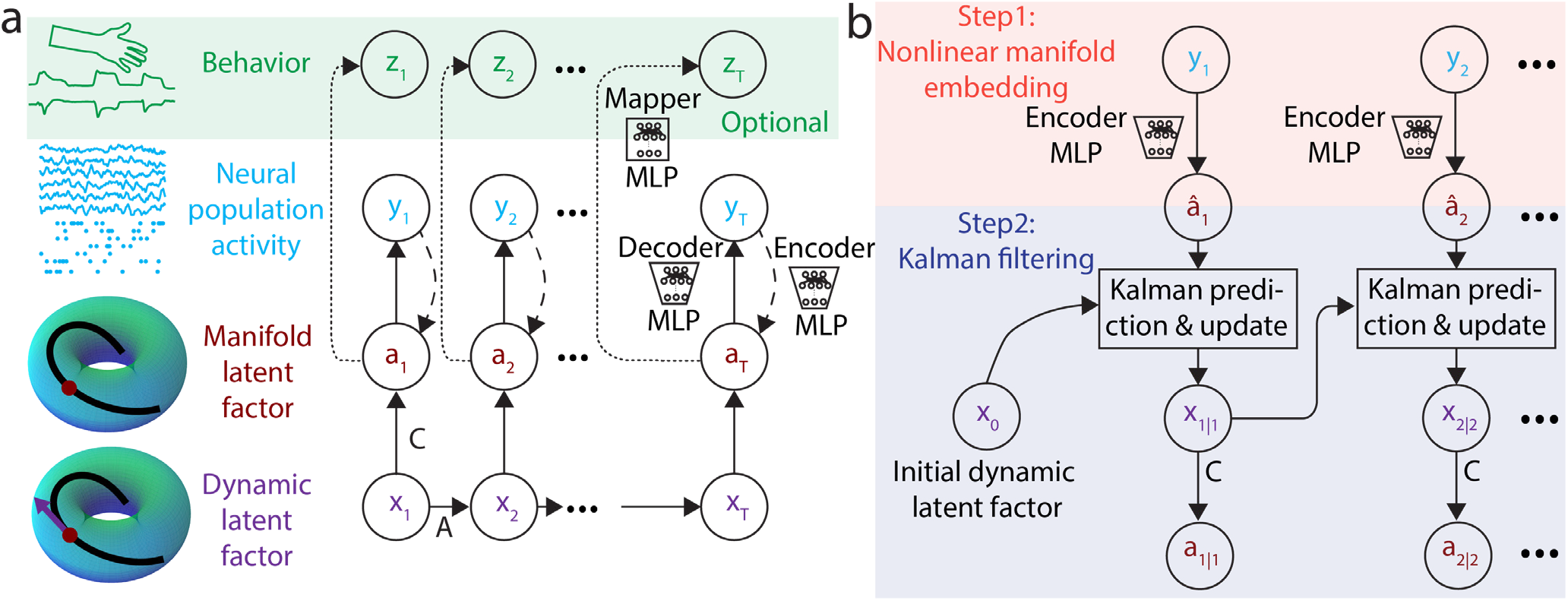
DFINE graphical model and its flexible inference method. **(a)** The DFINE model with two sets of latent factors is shown. These sets consist of the dynamic and manifold latent factors, respectively, which are separated to enable flexible inference while modeling nonlinearity. The relationship between the manifold latent factors and neural observations is modeled with an autoencoder with MLP encoder and decoder networks, where the manifold latent factor is the bottleneck representation. The dashed line from neural observations to the manifold latent factor is only used for inference and is not part of the generative model. The dynamic and manifold latent factors together form an LDM with the manifold factors being noisy observations of the dynamic factors, which constitute the LDM states. The temporal evolution of the dynamic latent factors is described with a linear dynamic equation. All model parameters (LDM, autoencoder) are learned jointly in a single optimization by minimizing the prediction error of future neural observations from their past. In the unsupervised version, after training the DFINE model, we use a mapper MLP network to learn the mapping between the manifold latent factors and behavior variables. We also extend to supervised DFINE where the mapper MLP network is simultaneously trained with all other model parameters in an optimization that now minimizes both neural and behavior prediction errors (**Methods**). **(b)** The inference procedure with DFINE is shown. We first get a noisy estimate of manifold latent factors using the nonlinear manifold embedding at every time point. With the aid of the dynamic equation, we use Kalman filtering to infer the dynamic latent factors **x**_*t*|*k*_ and refine our estimate of the manifold latent factors **a**_*t*|*k*_, with subscript *t*|*k* denoting inference at time *t* from all neural observations up to time *k*, **y**_1:*k*_. In addition to real-time filtering (*t*|*t*) which is displayed, DFINE can also perform smoothing or filtering in the presence of missing observations (**Methods**). Inference method is the same for unsupervised and supervised DFINE and is done purely based on neural observations as shown here (only model training is different) (**Methods**).

Further, inference is done recursively – such that the current inferred factor can be used to get the next inferred factor without the need to reprocess the neural data –, thus enabling computational efficiency and real-time implementation.

The manifold latent factors are taken as a lower-dimensional representation of the neural population activity, and the mapping between the two is characterized with an autoencoder whose decoder and encoder networks are modeled by multi-layer perceptrons (MLP) (**Fig. 1a**). We use MLPs to model nonlinearities because they are universal approximators of any nonlinear function under mild conditions^40^. Having captured the nonlinearity with the autoencoder, we now enable the model to have flexible inference properties by having the dynamic and manifold latent factors form an LDM (**Fig. 1a**). In this LDM, the manifold latent factors are noisy observations from the dynamic latent factors that constitute the LDM states and whose time-evolution is described through a linear dynamic model with additive noise (**Fig. 1a**). Using backpropagation, we jointly learn all the model parameters by minimizing the prediction error of future neural observations from past neural observations, measured using the root mean-squared error (RMSE). Since both dynamic and manifold latent factors are learned together in an end-to-end gradient-descent optimization, DFINE learns the best nonlinear manifold over which dynamics can be approximated as linearly as possible (**Discussion**).

For situations when specific behavioral variables are of interest and available during training, we extend DFINE to supervised DFINE so that learning of the model is informed by how predictive the learned manifold latent factors would be not only of future neural observations but also of behavioral variables. This is done during training by introducing an extra link from the manifold latent factors to continuous behavior variables (**Fig. 1a**) – termed the mapper network modeled with an MLP –, and by modifying the cost function to include both behavior and neural prediction errors (**Methods**). This additional link is purely added during training and removed afterwards during inference on test data. This leads to a learned model that is identical to the original model in terms of architecture and inference but just with different parameter values. Importantly, inference is again done purely from neural observations (**Fig. 1b** and **Methods**).

In addition to showing that DFINE enables the new capability of combining nonlinear neural network modeling with flexible inference, we also compare it to benchmarks of linear LDM and nonlinear SAE. To show the generalizability of DFINE across behavioral tasks, brain regions and neural signal types, we perform our analyses across multiple independent datasets. For SAE, we use the architecture named latent factor analysis via dynamical systems or LFADS^4^, which is a common benchmark nonlinear model of neural population activity^12, 29, 32^. For each dataset and algorithm, we infer the latent factors from the trained models. The latent factors correspond to the manifold latent factors in DFINE (**Methods**), to the state in the state-space model in LDM^21^, and to the dynamic factors (the representation layer right after the generator RNN) in SAE^4^. Unless otherwise stated, we use smoothing to infer the latent factors for all methods. We report all the quantifications using five-fold cross-validation, where the values are calculated in the held-out test set (see **Methods**). After extracting the latent factors, in the training set, we learn classification or regression models for discrete or continuous behavioral variables, respectively.

We quantify the cross-validated behavior prediction accuracy with area under the curve (AUC) of the receiver operating characteristic^41^ for discrete classifiers and with Pearson’s correlation coefficient (CC) for continuous regressions. The neural prediction accuracy is calculated with one-step-ahead prediction accuracy, the accuracy of predicting neural observations one step into the future from their past (see **Methods**). We also evaluate the neural reconstruction accuracy, defined as how well inferred latent factors – whether via causal filtering or via non-causal smoothing – reconstruct the current neural observations. Error values are computed in normalized RMSE (NRMSE) defined as RMSE normalized by the variance of the ground-truth observations to allow pooling the values across sessions given scaling differences (**Methods**). To assess how well the structure of the manifold is revealed in single-trials, we apply topological data analysis^42^ (TDA) on the extracted latent factor trajectories in test sets (**Methods**).

### Neural recordings and experimental tasks

We demonstrate the DFINE method using both extensive numerical simulations as well as diverse datasets containing distinct behavioral tasks, brain regions, and neural signal types to show the generalizability of the method. We use datasets containing four behavioral tasks as follows.

#### Saccade dataset

A macaque monkey (Monkey A) performed saccadic eye movements toward 1 of 8 peripheral targets on a screen in a visually-guided oculomotor delayed response task, which we refer to as the saccade task in short^43^ (**Fig. 2a**, **Methods**). Each trial started by presenting a central fixation square to which the monkey was required to maintain fixation, followed by Target On, Go, Saccade Start and End time events (**Fig. 2a**, **Methods**). We define the preparation period as the time between Target On and Go events, and the movement period as the time between Go and End events (**Fig. 2a**). We processed raw LFP signals in lateral prefrontal cortex (PFC) given their importance in saccadic eye movement representation^12, 44^ (**Methods**).

**Figure 2.**
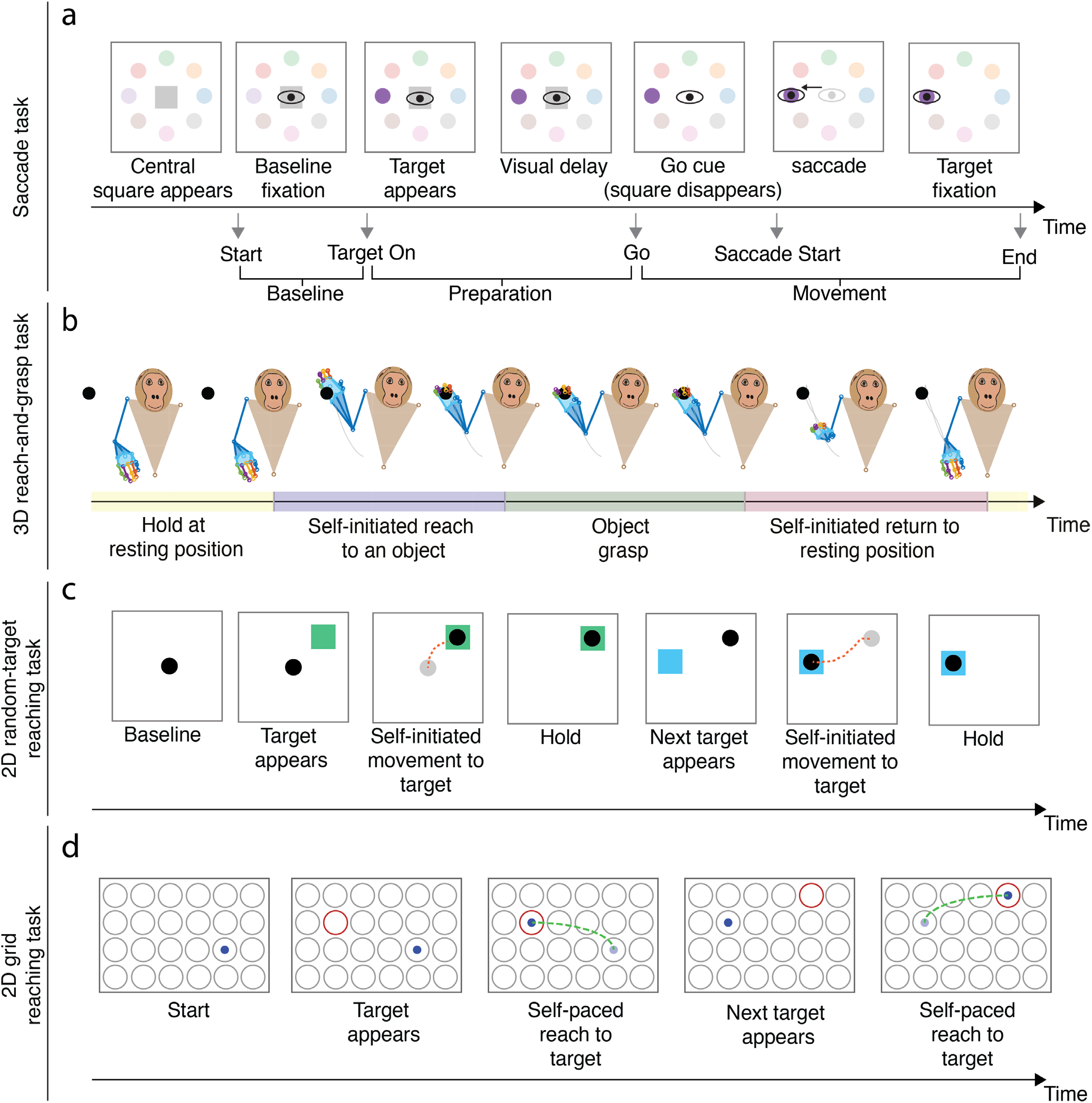
Experimental tasks. **(a)** Events for the saccade task are shown. At the beginning of each trial, the subject is required to fixate its eyes on a baseline location (gray square). After fixation, the target is illuminated on the screen (Target On event). After a visual delay, the fixation square disappears, which signals a go command (Go event). Then the subject performs the saccade (Saccade Start event) and holds fixation on target to receive a liquid reward (End event). We define the preparation and movement periods as the durations between Target On and Go events, and the duration between Go and End events, respectively. **(b)** In the naturalistic 3D reach-and-grasp task, an experimenter continuously moved a wand to diverse locations within a 3D area in front of the subject. The subject naturalistically reached to the object on the wand, grasped it, and returned its arm to the resting position. Movements were self-initiated without a specific go cue or timing instructions to isolate any parts of movements. **(c)** In the 2D random-target reaching task, the subject controlled a cursor (black circle) using a manipulandum. The task consisted of several sections, with each section having 4 sequential reaches to random targets (shown by colored squares) that appeared on the screen (only 2 reaches are shown here for simplicity). The subject performed self-initiated movements towards the targets. After a brief hold on a target, the next random target appeared at a pseudo-random location on the screen. **(d)** In the 2D grid reaching task, the subject controlled a cursor (blue dot) on a 2D grid in a virtual reality environment by moving its fingertip. Once the target (red circle) appeared on one of the circular locations on the grid (gray circles), the subject performed a self-paced movement towards the target, after which another target appeared from the set of possible targets.

#### Motor datasets

We used 3 independent motor datasets to show generalizability. For all motor datasets, we took the Gaussian smoothed spike counts as the neural signal to be modeled (**Methods**). The first motor dataset was a 3D naturalistic reach-and-grasp task, where the monkey performed naturalistic reach-and-grasps toward diverse locations in 3D space while the 3D endpoint hand position and velocity were measured and taken as the behavior variables^11^ (Monkey J, **Fig. 2b**, **Methods**). The neural recordings covered primary motor cortex (M1), dorsal premotor cortex (PMd), ventral premotor cortex (PMv) and PFC. The second motor dataset was a publicly available 2D random-target reaching task^45, 46^, where PMd activity was recorded while the monkey made sequential 2D reaches on a screen using a cursor controlled with a manipulandum, and the 2D cursor position and velocity were tracked as the behavior (Monkey T, **Fig. 2c**, **Methods**). The third motor dataset was a publicly available 2D grid reaching task^47, 48^, where M1 activity was recorded while the monkey controlled a cursor on a 2D surface in a virtual reality environment via its fingertip movements whose 2D position and velocity were tracked as the behavior (Monkey I, **Fig. 2d**, **Methods**).

### DFINE successfully learns the dynamics on diverse nonlinear manifolds and enables flexible inference in simulated datasets

We first validated the DFINE model and its learning algorithm in numerical simulations. Given the plausibility of ring-like, spiral-like and toroidal structures in neural population activity in prior studies^6, 10, 15, 49^, and to show the generality of the method to manifold types, we simulated trajectories on Swiss roll, Torus and ring-like manifolds as a proof of concept (**Fig. 3**, **Methods**). We synthesized 30 different simulated sessions (each with 250 trials) with randomly selected noise values and with the manifolds uniformly chosen from the above possibilities (**Methods**). Given the noisy nature of neural recordings, the simulation observations were taken as the noisy realizations of the trajectories on the manifold (**Fig. 3b**).

**Figure 3.**
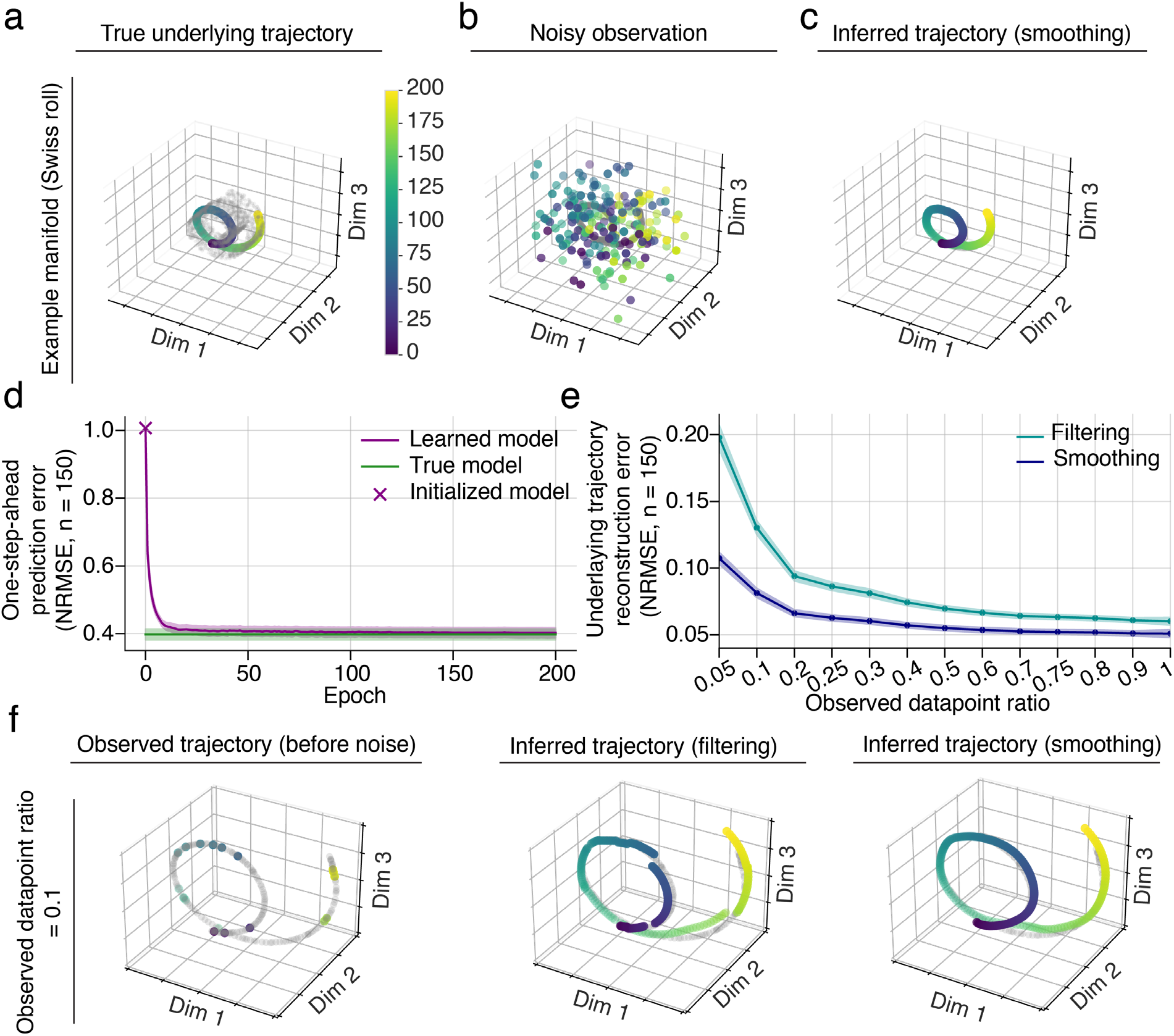
DFINE successfully learns the dynamics on nonlinear manifolds and enables inference in the presence of missing observations in simulated datasets. **(a)** A sample simulated trajectory is shown for an example manifold (Swiss roll). The gray points are samples from the underlying manifold. Color evolution on the trajectory indicates time evolution into a trial as shown in the color bar. **(b)** Noisy observations of the trajectory are shown with the same color convention as in (a) and on top of the gray true trajectory. **(c)** After learning the DFINE model, inferred trajectory with smoothing is shown, which is essentially on top of the true trajectory represented with gray dots and masking it. **(d)** The learned models’ one-step-ahead prediction error converges to that of the true models. The plot shows the mean of one-step-ahead prediction error for the learned and true models across all simulated sessions, cross- validation folds and trials. The shaded areas represent the 95% confidence bound of the mean. DFINE starts from a randomly initialized model, which has a chance-level one-step-ahead-prediction of 1. **(e)** Neural reconstruction error between the inferred and true trajectories is shown for smoothing and filtering across various observed datapoint ratios. Dots represent the mean across simulated sessions and cross-validation folds and shaded area represents the 95% confidence bound. Error is essentially unchanged as samples are dropped in a large range of observed datapoint ratios and is significantly lower than chance-level of 1 for all ratios (*P* < 8.7 × 10^−7^, one-sided Wilcoxon signed-rank test). Smoothing is more accurate than filtering across all ratios (*P* < 8.7 × 10^−7^, one-sided Wilcoxon signed- rank test). **(f)** An example trajectory with missing observations and its smoothing and filtering inference for observed datapoint ratio of 0.1. Colored dots in the left panel are observed datapoints and the gray dots are shown to visualize the true underlying trajectory.

We found that DFINE can correctly infer the trajectories on the manifolds, even from noisy observations (**Figs. 3a**-**c**) and even in the presence of missing observations (**Fig. 3f**). First, the learned model’s one-step-ahead prediction error converged to that of the true model (**Fig. 3d**). Indeed, the difference between learned and the true model errors, normalized by the true model error, decreased from 1.564 ± 0.110 (mean ± sem) for randomly initialized models to 0.012 ± 0.004 for the learned model, indicating convergence (**Fig. 3d**). Second, the same DFINE model enabled inference in the presence of missing observations (**Figs. 3e-f**). To show this, we randomly dropped neural observation datapoints from each trial to achieve a desired observed datapoint ratio, which is defined as the ratio of the datapoints that are maintained/not-dropped to the total number of datapoints (see **Methods**). DFINE predictions even for ratios as low as 0.2 were similar to when all datapoints were retained, showing that DFINE could use the learned dynamics to compensate for missing observations (**Fig. 3e**). Further, even for ratios as low as 0.05, DFINE still performed better than chance-level of 1 (**Fig. 3e**). Third, smoothing significantly improved the inference of trajectories because it could also leverage the future neural observations, and this improvement due to smoothing was more prominent in the lower observed datapoint ratios (**Fig. 3e**). Indeed, smoothing led to successful predictions even for ratios as low as 0.1 (**Fig. 3f**). Overall, the simulation analysis showed that DFINE can learn the dynamics on diverse nonlinear manifolds, perform flexible inference both causally (filtering) and non-causally (smoothing), and succeed even in the presence of missing observations.

### DFINE extracts single-trial latent factors that accurately predict neural activity and behavior

We then applied DFINE on the four independent datasets to show that it not only allows for flexible inference but also for accurate neural and behavior prediction. We compared DFINE’s single-trial latent factors to those of the linear (LDM) and nonlinear (SAE) benchmarks (**Methods**). We first visualized the condition-average and single-trial latent factor trajectories for the saccade task during both preparation (**Fig. 4a**) and movement periods (**Fig. 4b**), where condition is defined as the saccade target. We found that the latent factors inferred with DFINE not only captured inter-condition variabilities during movements even in single-trials (**Fig. 4b**), but also exhibited smooth trajectories with a discernable manifold structure in these noisy single-trials (**Figs. 4a** and **4b**). This was unlike LDM that generally had noisier single-trial latent factor trajectories, and unlike SAE whose latent factor trajectories, while smooth, captured smaller inter-condition variabilities in single-trials (**Figs. 4a** and **4b**).

**Figure 4.**
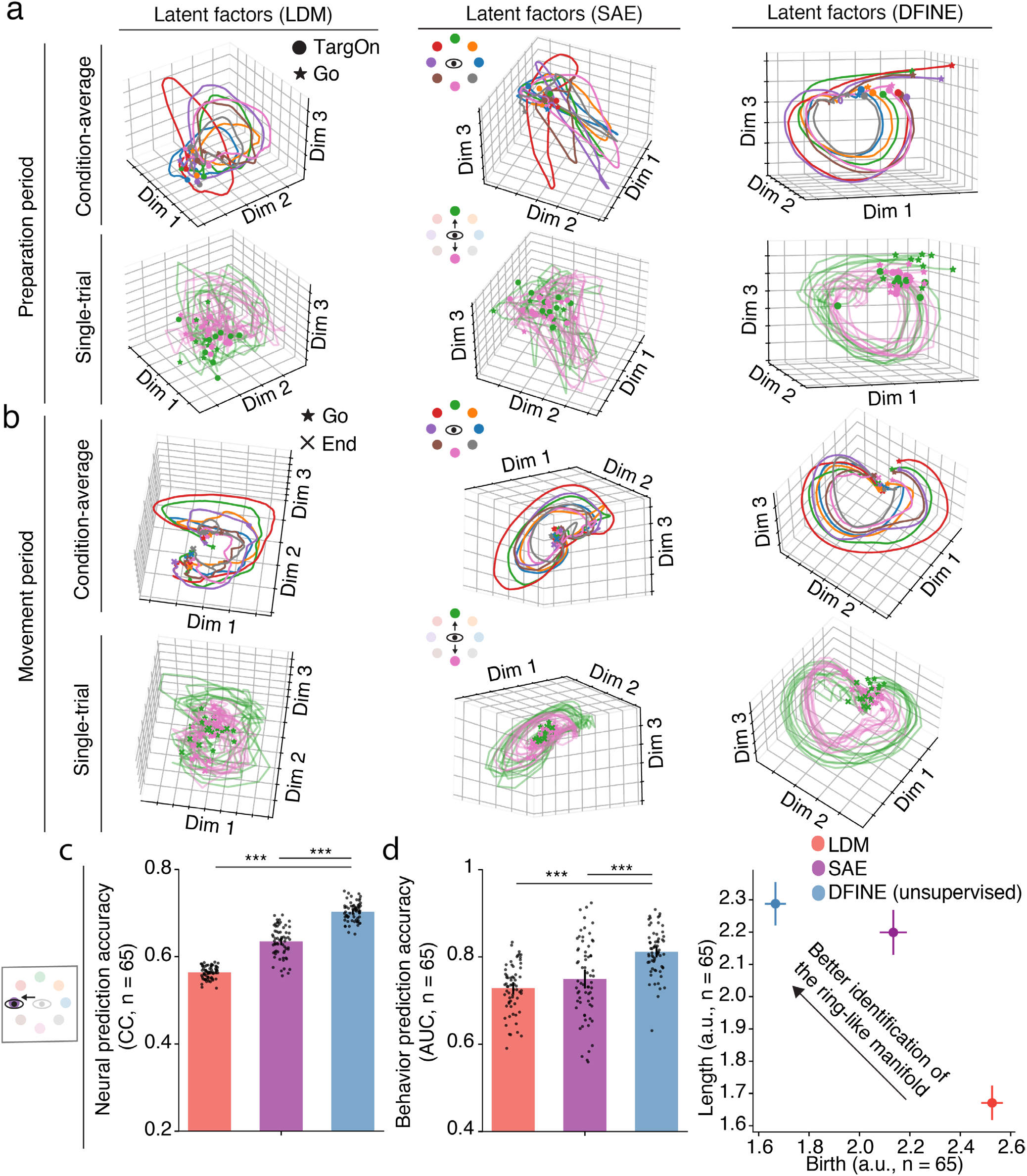
In the saccade dataset, DFINE outperformed benchmark methods in behavior and neural prediction accuracy, and more robustly extracted the ring-like manifold in single-trials. **(a)** Condition-average (top row) and single-trial (bottom row) latent factor trajectories for an example session are shown for DFINE, LDM and SAE during the preparation period. Each color represents one target, i.e., condition. **(b)** Similar to (a), for the movement period. **(c)** DFINE had significantly higher neural prediction accuracy compared to LDM and SAE. All models had 16-dimensional latent factors (see **Supplementary Fig. 1** for convergence). Black dots represent the accuracy in each cross-validation fold of each session, bars represent the mean, and error bars represent the 95% confidence bound. Asterisks indicate significance of comparison (*: *P* < 0.05, **: *P* < 0.005 and ***: *P* < 0.0005, one- sided Wilcoxon signed-rank test). **(d)** Behavior prediction measured by target classification accuracy was better with DFINE compared to LDM and SAE. Convention is the same as in (c). **(e)** TDA analysis on single-trial latent factors during the movement period is shown. The top left area of the plots corresponds to more robust extraction of the ring-like manifold (**Methods**). Circles represent mean across sessions and cross-validation folds and error bars represent the 95% confidence bound. TDA’s most persistent 1-dimensional hole had a significantly earlier birth and lasted significantly longer for DFINE compared to LDM and SAE (*P* < 5 × 10^−4^, one-sided Wilcoxon signed-rank test).

We next quantified this ability to capture inter-condition variability by computing the single-trial neural and behavior prediction accuracies of 16-dimensional (16D) latent factors for each method. We picked this dimension because it was sufficient for the performance of all methods to converge across all datasets (**Supplementary Fig. 1**; SAE’s dynamic state dimension is taken to be much higher at 64, see **Discussion** and **Methods**). Note that during a given trial, SAE does not predict the neural observations into the future from its past because it needs to use *all* neural observations until the end of a trial to get the initial condition and then to extract the factor trajectories at every time-step from this initial condition. Thus, while we computed the one-step-ahead neural prediction error for DFINE and LDM using only past neural observations, we gave SAE the advantage of doing neural reconstruction with smoothing based on both past and future neural observations instead.

Despite this advantage given to SAE, DFINE was better at neural prediction not only compared with LDM but also compared with SAE across all datasets (**Fig. 4c**, **Figs. 5a**, **5d** and **5g**). In comparison with SAE and LDM, respectively, DFINE improved the accuracy of neural prediction in the 3D naturalistic reach-and-grasp task by 19.9 ± 1.8% and 49.0 ± 3.7% (**Fig. 5a**), in the 2D random-target reaching task by 56.7 ± 26.7% and 43.9 ± 7.3% (**Fig. 5d**), in the 2D grid reaching task by 27.8 ± 6.5% and 25.9 ± 2.2% (**Fig. 5g**), and in the saccade task by 10.9 ± 1.7% and 24.7 ± 1.3% (**Fig. 4c**). Similar results held for the comparison of DFINE with SAE in terms of neural reconstruction with smoothing (**Supplementary Fig. 2**, see also **Discussion**).

**Figure 5.**
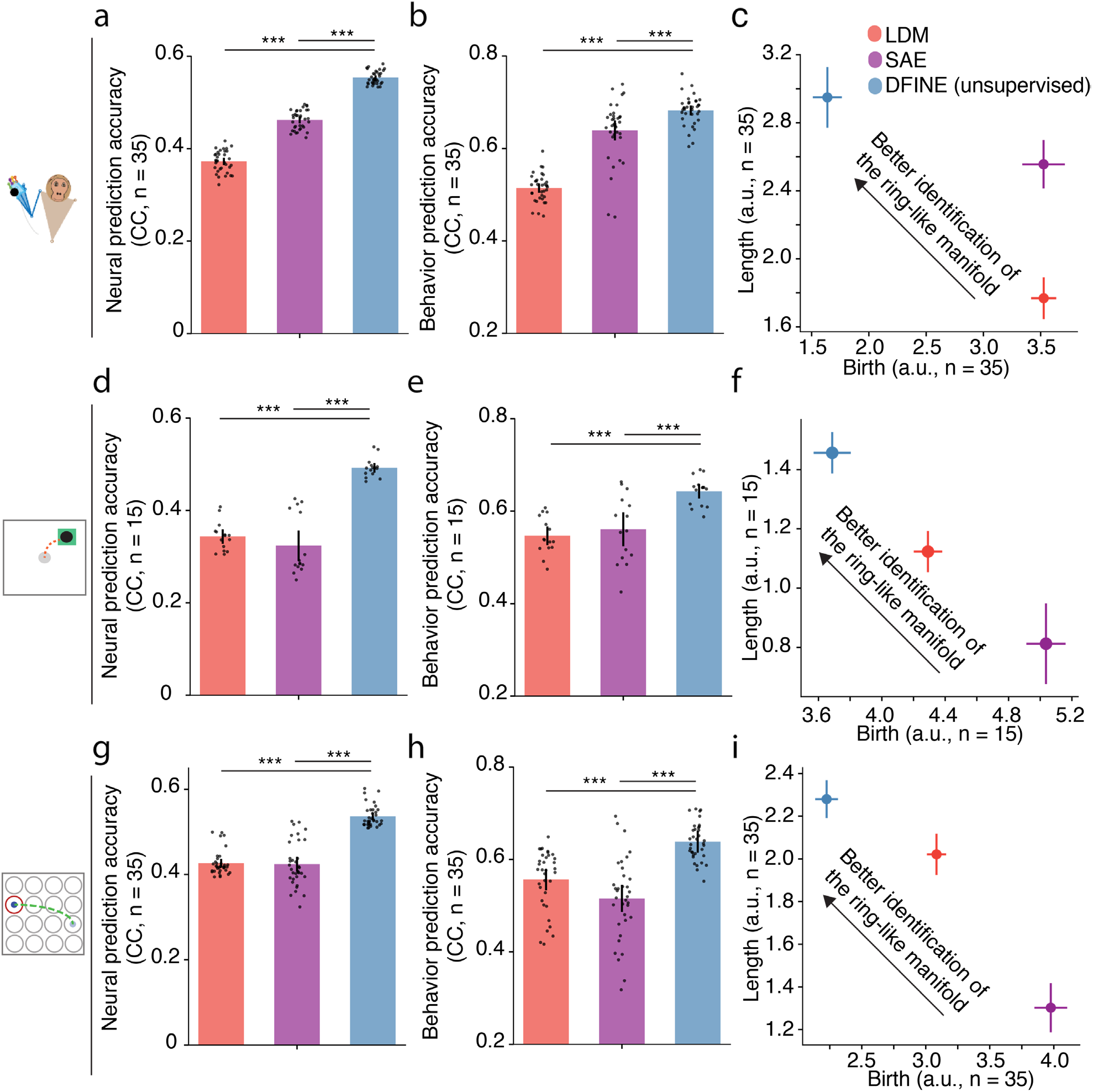
In the motor datasets, DFINE outperforms benchmark methods in behavior and neural prediction accuracy, and more robustly identifies the ring-like manifold structure in single-trials. Figure convention for bars, asterisks for significance and for the TDA plots are the same as Fig. 4. DFINE again outperformed benchmarks in terms of neural prediction, behavior prediction, and robust extraction of the manifold structure in the naturalistic reach-and-grasp task **(a-c)**, random-target reaching task **(d-f)**, and 2D grid reaching task **(g-i)**.

In addition to its better neural prediction, DFINE also had higher behavior prediction accuracy compared to LDM and SAE in all tasks (**Methods**). In the motor tasks, improvements compared to SAE and LDM, respectively, were 7.7 ± 5.7% and 33.0 ± 4.0% in the 3D naturalistic reach-and-grasp task (**Fig. 5b**), 16.8 ± 17.1% and 17.9 ± 3.9% in the 2D random-target reaching task (**Fig. 5e**), and 21.2 ± 7.5% and 11.6 ± 7.0% in the 2D grid reaching task (**Fig. 5h**). Also, for the saccade task, DFINE latent factors better predicted the saccade target class during the movement periods (**Methods**), with the saccade target classification AUC being 9.7 ± 3.9% and 11.8 ± 2.9% better than SAE and LDM, respectively (**Fig. 4d**).

These results demonstrate that in addition to enabling flexible inference, DFINE’s single-trial latent factors were more predictive of both behavior and neural activity compared to the benchmark linear and nonlinear methods.

### DFINE more robustly extracts the manifold structure in single-trials

Consistent with its better behavior and neural prediction, DFINE also more robustly captured the nonlinear manifold structure in single-trial data compared with both LDM and SAE. First, visualization of DFINE revealed a ring-like manifold structure in both condition-average and single-trial latent factors during both preparation and movement periods in the saccade task (**Figs. 4a** and **4b**) and during the movement periods in the motor tasks (**Supplementary Fig. 3**). We also observed that this ring-like structure was much less apparent in single trials for LDM and SAE (e.g., **Figs. 4a**). To quantify this observation and whether DFINE was better able to extract this manifold in single-trials, we used TDA which uses persistent homology to quantify whether there exist holes in data, and if so how many^42^ (**Methods**). TDA finds multi-dimensional holes (e.g., 1D hole is a ring and 2D hole is a 2D void) in data by growing the radius of ***ϵ***-balls around datapoints (**Methods**). If holes exist, a model that finds holes which are born earlier and last longer – i.e., are more persistent – is more robust in revealing the manifold structure in single-trials. Consistent with observing a ring in low-dimensional visualizations above, TDA revealed a persistent 1D hole in low-dimensional latent factors, which we then analyzed.

Compared with benchmarks, in DFINE, the birth of the TDA’s most persistent 1D hole was significantly earlier and its length was significantly larger during both preparation and movement periods for the saccade task (**Fig. 4e** and **Supplementary Fig. 4**; *P* < 5 × 10^−4^, one-sided Wilcoxon signed-rank test), and during movement periods for all motor tasks (**Figs. 5c, 5f, 5i**; *P* < 5 × 10^−4^, one-sided Wilcoxon signed-rank test). These results show that the ring-like manifold structure was more robustly captured with DFINE than both LDM and SAE.

### DFINE can also be extended to enable supervision and improve behavior prediction accuracy

So far, we presented the results of unsupervised DFINE in which the model is trained unsupervised with respect to behavior and to optimize neural prediction alone. To allow for considering continuous behavior measurements when available during training, we next developed supervised DFINE to train a model with latent factors that are optimized for both neural and behavior predictions (**Fig. 1a**; **Methods**). To validate the supervised DFINE method, we applied it to the motor datasets in which continuous behavior measurements were available during training.

We found that supervised DFINE successfully improves the behavior prediction accuracy even though its model and inference architectures are identical to those in the unsupervised DFINE, and even though its inference only takes the same neural observations as input (**Fig. 6**; **Methods**). The latent factors of supervised DFINE better distinguished different task conditions across all motor datasets compared to those of unsupervised DFINE (**Figs. 6a**, **6c** and **6e**). Consistent with this observation, supervised DFINE significantly improved the behavior prediction accuracy compared to unsupervised DFINE across all motor datasets (**Figs. 6b**, **6d** and **6f**). These improvements were 13.6 ± 2.7% in the 3D naturalistic reach-and-grasp task (**Figs. 6b**), 22.3 ± 1.8% in the 2D random-target reaching task (**Figs. 6d**), and 24.5 ± 2.5% for the 2D grid reaching task (**Figs. 6f**). Also, as expected, the neural prediction accuracy of supervised DFINE was significantly lower than unsupervised DFINE across all datasets (**Supplementary Fig. 5)**; this is because supervised DFINE’s latent factors are optimized for both behavior and neural prediction while those of unsupervised DFINE are optimized purely for neural prediction.

**Figure 6.**
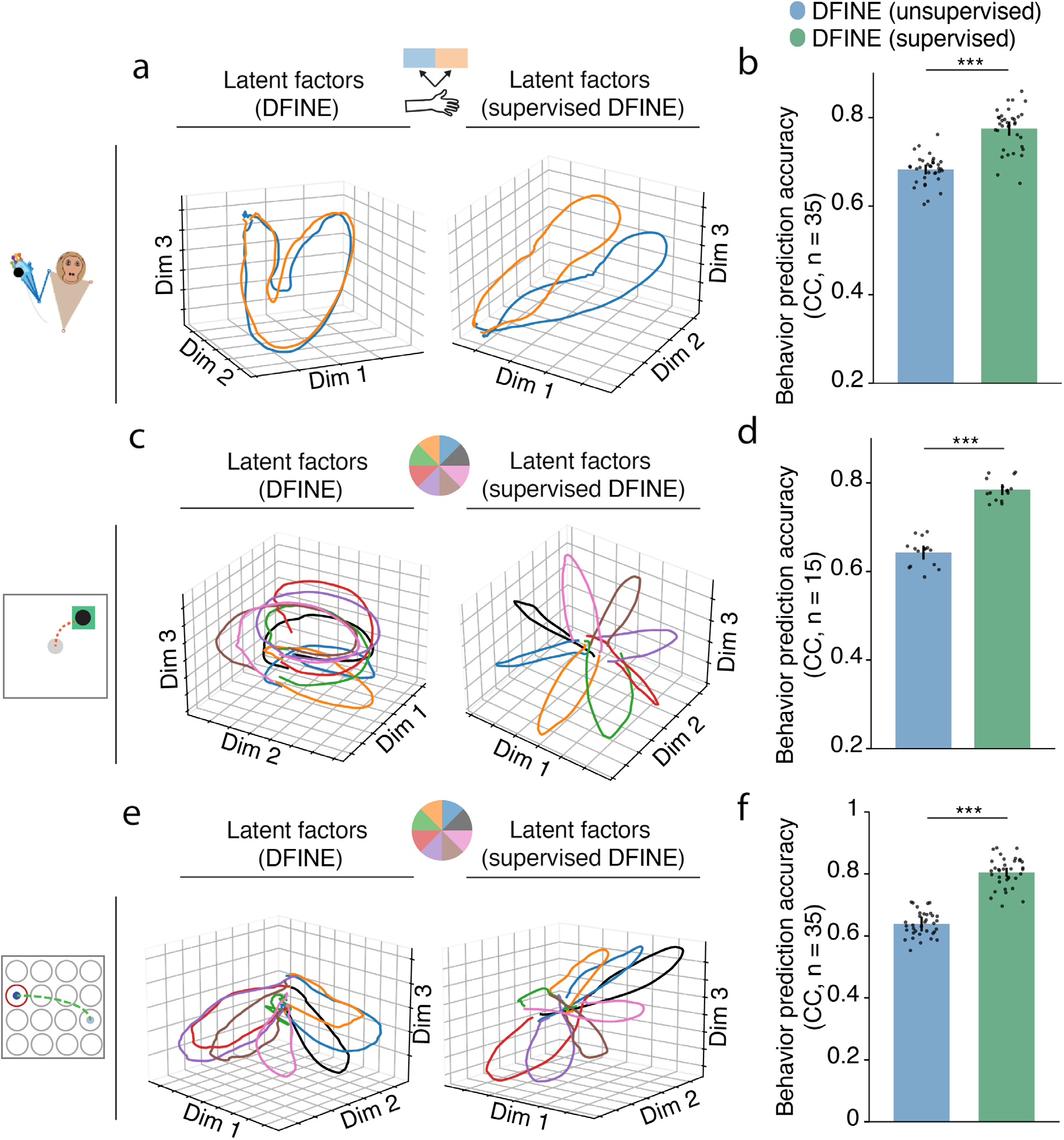
Supervised DFINE extracts latent factors that are more behavior predictive. **(a)** Examples of condition-average latent factor trajectories for the unsupervised (left) and supervised (right) DFINE are shown for the 3D reach-and-grasp task. Each color represents one condition (i.e., movement to left or right; **Methods**). Supervised DFINE better separates different conditions. **(b)** Supervised DFINE improved the prediction of behavior, i.e., continuous position and velocity, in the 3D reach-and-grasp task. Dots represent cross-validation folds across experimental sessions and the convention for bars, error bars and asterisks are the same as Fig. 4. Similar results held for the 2D random target reaching task **(c-d)** and 2D grid reaching task **(e-f)**, where here each condition is a direction angle interval/sector (for example, all movements whose direction angle is between 0-45 degrees regardless of where they start/end; **Methods**).

### DFINE can perform flexible inference with missing observations

We next investigated whether DFINE can perform inference even in the presence of missing neural observations (**Fig. 7**). To do so, we uniformly dropped neural observations throughout the recordings (**Methods**) and performed inference in two ways: 1) filtering inference (causal) that only uses the past and present available neural observations, 2) smoothing inference (non-causal) that uses all available neural observations.

**Figure 7.**
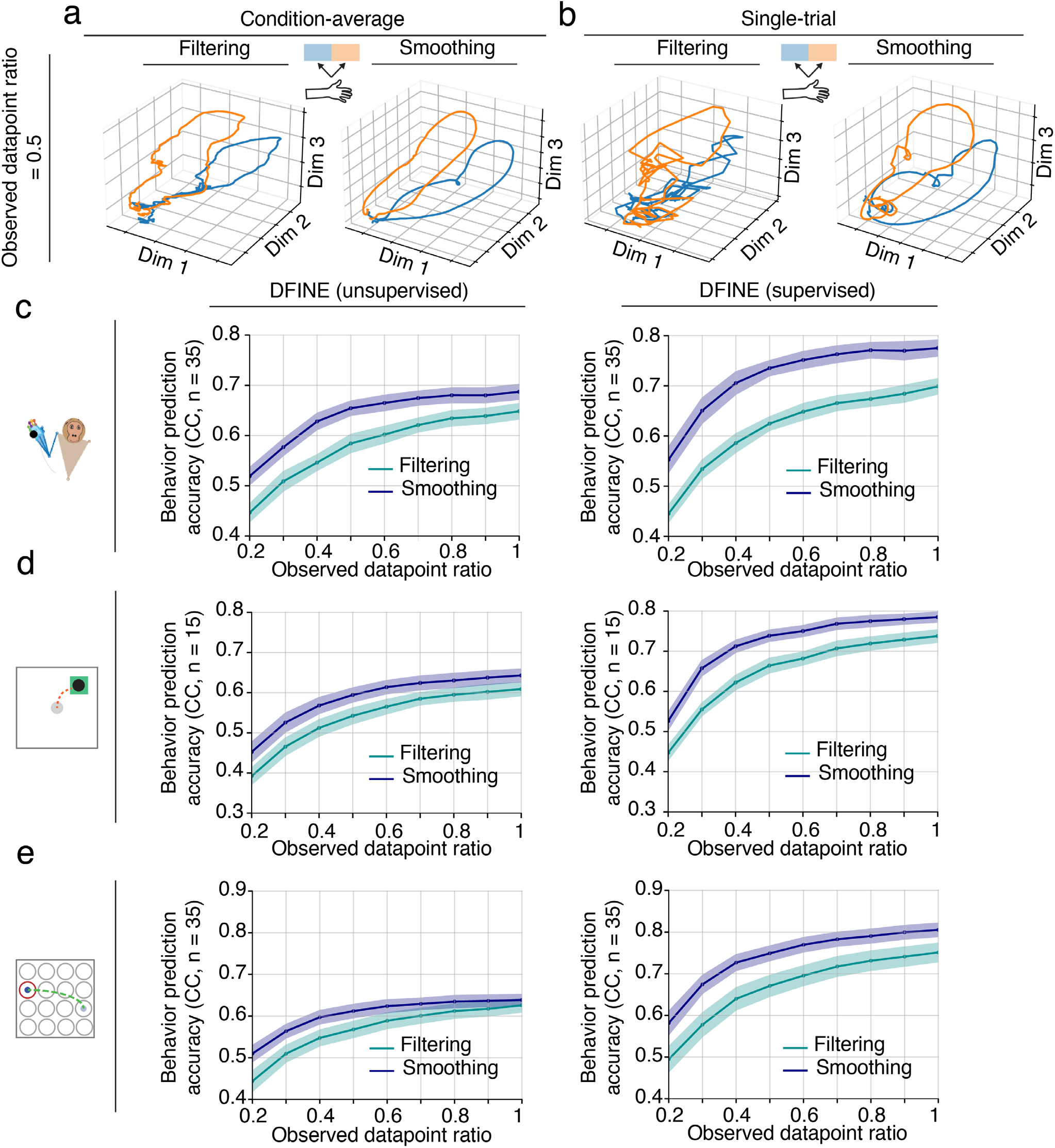
DFINE can perform both causal and non-causal inference with missing observations and do so more accurately through non-causal inference. (a-b) Examples of condition-average (a) and single-trial (b) latent factor trajectories for filtering and smoothing inference with missing observations in the 3D reach-and-grasp task. Both DFINE filtering and smoothing captured the low-dimensional structure in single-trials, with smoothing doing so more accurately than filtering. **(c)** Behavior prediction accuracies of filtering and smoothing inferences are shown across various observed datapoint ratios for the 3D naturalistic reach-and-grasp task. Lines represent the mean across experimental sessions and cross-validation folds, and the shaded areas represent the 95% confidence bound. **(d)** Similar to (c), for the 2D random-target reaching task. **(e)** Similar to (c), for the 2D grid reaching task. See also **Supplementary** Fig. 6.

We found that behavior prediction accuracies of DFINE remained relatively unchanged even when dropping up to 40% of observations (0.6 ratio), and remained well above the chance-level of 0 even when dropping 80% of observations (lowest 0.2 ratio) (**Figs. 7c**-**e**; *P* < 5 × 10^−4^, one-sided Wilcoxon signed-rank test). Also, behavior prediction accuracy with smoothing inference was significantly better than that with filtering inference across all observed datapoint ratios (**Figs. 7c**-**e**). **Figs. 7a** and **7b** visually demonstrate that the smoothing inference yields more accurate reconstruction of the low- dimensional latent trajectories as it leverages both past and future available observations. Indeed, the smoothed trajectories at observed datapoint ratio of 0.5 (**Fig. 7a**) look very similar to those for observed datapoint ratio of 1 (**Fig. 6a right panel**), showing ability to compensate for missing observations.

We then compared DFINE with SAE in terms of inference in the presence of missing observations. Because SAE’s decoder network is modeled with an RNN, it is structured to take neural observation inputs at every time-step. To handle missing observations for SAE at inference, we imputed them with zeros in the test set as previously done^33, 34^, extracted the latent factors, and computed the associated behavior prediction accuracy. DFINE outperformed SAE and, interestingly, this improvement grew larger at lower observed datapoint ratios (**Supplementary Fig. 6**). Indeed, DFINE’s degradation in performance due to missing observations was much lower than that of SAE (**Supplementary Figs. 6b**, **6d** and **6f**). These analyses show that DFINE can flexibly compensate for missing observations and perform both causal and non-causal inference with missing data. These analyses also show that DFINE’s non-causal inference can leverage future data for more accurate prediction when real-time processing is not needed.

## Discussion

We developed DFINE, a new nonlinear neural network model of neural population activity that enables the new capability for flexible inference, whether causally or recursively in real-time, non-causally to leverage the entire data length, or in the presence of missing neural observations. This flexible inference capability is critical both for future neurotechnology and to study the neural basis of behavior in causal experiments. In addition to enabling this new capability for flexible inference, DFINE more accurately predicted both the neural and behavioral data and more robustly discovered the low-dimensional neural manifold compared with linear and nonlinear benchmarks. We also developed a new algorithm to allow supervision for DFINE and to extract latent factors that are more behavior predictive, with no changes to the model architecture and inference properties otherwise. These capabilities and advantages generalized across four independent datasets with different behavioral tasks, brain regions, and neural signal modalities.

### DFINE allows for neural network modeling while also enabling flexible inference

Many studies have shown that neural population activity can be summarized with a significantly lower-dimensional latent manifold structure^1–12, 14, 50–52^, often using linear dimensionality reduction methods^1–3, 5, 7–12, 14, 50, 52^. To model the nonlinearities in neural population activity and better learn the underlying manifold structure^4–6, 10, 29–32, 37, 38, 51, 53–62^, while some studies have used spline loop fitting^6^ or extensions of isometric feature mapping (Isomap)^51^, most works have leveraged the power of neural networks^4, 5, 10, 29–32, 37, 38, 53–62^. As models of neural data, neural networks have either been in the form of generative models^4, 13, 29–32, 61^, whether static^61^ or dynamic^4, 13, 29–32^, or in the form of predictive models or decoders^37, 38, 60^. DFINE provides a dynamic generative model of neural data but with the major difference that it provides the new capability for flexible inference, in addition to being accurate in neural, behavior, and manifold prediction. DFINE has flexible inference in that it simultaneously enables recursive real-time/causal inference, non-causal inference, and inference in the presence of missing observations. Such flexible inference is critical to enable both real-time neurotechnology and accurate testing of scientific hypotheses, but is not achieved by prior neural network models of population activity.

Prior generative network models do not provide these flexible inference properties. This is because their inference cannot be solved analytically and is usually performed with an inference/recognition network that is trained in conjunction with the generative network to approximate the posterior distribution of latent factors. Therefore, inference properties depend on how the inference network architecture is structured to process the neural observations. In many cases, the inference network structure does not allow for real-time recursive estimation of latent factors^4, 13, 29, 30, 32^, and/or does not directly enable inference in the presence of missing observations^4, 13, 30–32^. For example, SAEs^4^, which are often used as benchmarks for neural data, are suitable for non-causal processing and are not amenable to recursive causal/real-time inference. With every new neural observation, the encoder RNN has to update the generator RNN’s initial latent state at time zero; then the generator RNN needs to reset and re-estimate its latent states throughout all time-steps to infer the latent factor at the time of the new neural observation. This non-recursive procedure is a large burden for real-time inference as the current estimate of latent factors cannot be utilized to get the next time-step’s factors. Indeed, all the latent factors have to be re-computed for every single new neural observation. In contrast, DFINE allows for recursive inference without the need to re-compute any of the past or current factors. In terms of addressing missing observations, while imputation techniques such as zero-padding have been used for SAEs or in general for models with RNNs^33, 34^, it is known that such techniques usually yield sub- optimal performance^35, 36^. This is because imputing missing observations with zero in inference distorts the observation values. Similar to SAEs, in prior linear dynamical models with nonlinear embeddings^30^, inference again needs to use all neural observations, therefore recursive causal inference or inference with missing observations is not directly addressed. Similarly to these prior generative networks, prior predictive networks with RNN-based methods^37–39^ also do not allow for flexible inference because, this time, they do not enable non-causal inference given their forward RNN architecture and do not directly address missing observations as RNNs are structured to take inputs at every time-step. Further, unlike DFINE, these predictive networks do not directly learn a generative model of neural population activity and thus are largely used for decoding purposes rather than to study/infer neural population structure.

Beyond neuroscience, dynamical modeling of sequential data has great importance for other domains such as for processing of videos^63–65^ or text^66, 67^, with great progress made to date. However, neural data introduce distinct challenges that DFINE’s modeling architecture was specifically designed to handle. First, to consider the noisy nature of neural activity, we model and learn stochastic noise variables in DFINE. This allows us to model uncertainties and to fit the noise values to any specific dataset across diverse tasks, brain regions, and neural data modalities (e.g., spiking or LFP). In contrast, because video or text observations are deterministically observed or are much less noisy, in applications of videos and text, the stochastic noise variables are not necessarily included in the modeling or are tuned manually as hyperparameters^63–67^. In addition, instead of the common method of optimizing the evidence lower bound (ELBO) of data likelihood when noise variables exist^4, 30, 64^, we optimize the prediction accuracy of data into the future because optimizing ELBO could be challenging^68–71^ and a major goal of a dynamic model is future prediction; indeed, we observed instabilities when using ELBO as our optimization cost. Finally, unlike these video/text applications^63–67^, we also developed a supervised learning algorithm for DFINE to allow for modeling two different but related sets of sequential data (neural activity and behavior here) and motivate the latent factors to be predictive of not just one but both time-series/sequences. In addition to modeling of neural/behavior data in neuroscience, this supervised learning algorithm could show promise for future applications of video/text data where another measured time-series/sequence is also of interest, e.g., inferring latent factors of videos regarding movements^72^.

### Linear temporal dynamics on the nonlinear manifold and extension across neural modalities

To extend the linear dynamical models of neural population activity^1, 2, 11, 12, 21, 24–27^, several prior studies have built nonlinear models of temporal dynamics using neural networks^4, 29, 31, 32^ or Gaussian- process based methods^73–76^. While DFINE develops a nonlinear neural network model, it keeps the temporal dynamics on the manifold linear for several reasons. First, the separation of the dynamic and manifold latent factors and the linear form of the former is what allows for flexible inference. Second, since all manifold and dynamic model parameters are trained together in an end-to-end single optimization, DFINE learns the best nonlinear manifold over which dynamics can be approximated as linearly as possible. This joint training gives the model the flexibility to change the nonlinear manifold as needed such that the dynamics on top of it can be closely approximated as linear. Third, recent work that dissociates the source of nonlinearity in neural population activity has shown that linear dynamics in the presence of other nonlinearities such as embeddings may be sufficient in explaining the neural observations^39^. Consistent with this finding, our results suggest that the linear description of temporal dynamics on the nonlinear manifold in DFINE does not degrade the neural and behavior prediction as DFINE even outperforms fully nonlinear SAEs with nonlinear temporal dynamics. This is despite the fact that these SAEs can have even more general nonlinear dynamics than the time-varying dynamics in LDM extensions, such as switching LDMs^77–79^ (**Figs. 4** and **5**). Finally, DFINE can be extended to allow for locally linear dynamics as described below to better capture potential nonlinearities that may not be fully captured using the joint optimization of the nonlinear manifold factors and dynamic factors.

In the DFINE model, we used a Gaussian distribution for neural observations such that our architecture can generalize across various neural data modalities including not only spike counts but also field potentials, which are continuous-valued. Indeed, field potentials can provide a robust modality for neurotechnologies^17, 80^ and can have comparable performance to spiking activity when decoding certain behavior variables^11, 12, 44, 81–83^. Field potentials can also reveal larger-scale network computations during behavior for basic science investigations^11^. Beyond this, extending DFINE to support other observation distributions is a future direction.

DFINE’s major objective is to enable the new capability for flexible inference while also capturing nonlinearity, which is essential for neurotechnology. Nevertheless, to show that despite enabling this new capability, DFINE still allows for accurate modeling and inference, we compared it with SAEs using the LFADS architecture^4^ as a benchmark (**Methods**). For a fair comparison, similar to DFINE, we had LFADS use a Gaussian observation distribution to make it applicable to both spike counts and field potentials. To get a conservative estimate of the improvements in DFINE, we allowed LFADS to have a higher dynamics state dimension than DFINE (64 vs. 16; see **Methods**). For the choice of hyperparameters in LFADS, we picked the values given for one of the datasets in the original work^4^ that had the closest number of trials to our datasets (see **Methods**). It is possible that a comprehensive hyperparameter tuning can improve LFADS’s performance; however, this tuning would require significant training time given the complex architecture and its large number of hyperparameters. While an extension of LFADS, AutoLFADS^84^, could help with this tuning, the current AutoLFADS implementation does not support Gaussian observation distributions unlike LFADS and DFINE, and thus is not directly comparable to DFINE or applicable to the field potential modalities here. Further, AutoLFADS optimizes only the non-architectural hyperparameters during training and the architectural hyperparameter search (such as all layer/RNN latent state dimensions) still requires significant training resource/time. In contrast, DFINE uses the same architectural hyperparameters for all datasets in both its supervised and unsupervised versions, thus not requiring such an extensive search. This shows the simple and generalizable architecture and training of DFINE. Further, AutoLFADS is an SAE and so does not support flexible inference. Nevertheless, to test the choice of LFADS hyperparameters here, we applied LFADS with these hyperparameters to a publicly available spiking dataset during a maze task^85^ that has recently been tested with AutoLFADS. We found that LFADS with the chosen hyperparameters here had a behavior decoding performance similar to that reported for AutoLFADS on this same maze data^85^, suggesting the appropriateness of the chosen LFADS hyperparameters here.

### Revealing low-dimensional manifold structure in the saccade and motor datasets

Latent factor models have led to significant insight about neural population coding during various behavioral tasks. For example, rotatory low-dimensional structure in the neural population activity has been prevalently observed^1, 2, 4–6, 8–10, 14, 49, 55, 86, 87^, in the motor system during movements^1, 2, 4, 9, 10, 49, 55, 86, 87^ as well as other systems during tasks such as a syllables task^8^, in a ready-set-go task while performing a saccade^5^, in an exploration task in the head direction system^6^, and during auditory stimulation^14^. Here, DFINE found latent factors that consistently had ring-like manifold structures as revealed by visualization and TDA analysis. Importantly, DFINE extracted such ring-like manifold structures more robustly in single-trials compared with LDM and SAE in all datasets.

In addition to allowing for flexible inference, the separation of dynamic and manifold factors in DFINE can facilitate neuroscientific interpretation. We can interpret the manifold latent factors as capturing the global latent structure of the population code and the dynamic latent factors as revealing its local properties and variations. This is because the same manifold can be traversed in many ways, which are captured by the dynamic latent factors; thus, dynamic factors may correspond to local changes in behavior while manifold factors may relate to global changes in behavior. Combined with its accurate and robust discovery of the latent structure, this separation can also facilitate the use of DFINE for investigations and interpretations across diverse domains of neuroscience.

### Future directions

Given its flexible training using backpropagation, there are several ways to extend DFINE while maintaining all its current advantages such as flexible inference. Here, we modeled the output neural observations with a Gaussian distribution to be generalizable across various neural modalities whether spike counts or field potentials for example. Changing the output distribution to generalized linear models (GLM) such as Poisson/point process likelihood functions^11, 25–27, 30, 84, 88^ or multiscale observation models^11, 89–92^ can be explored in the future. Multiscale observation models require extra care to fuse information across multiple scales when recovering the latent manifold^93^. Finally, inputs could also be modeled in the future by incorporating them in the LDM part of the model for example for stimulation applications^17, 94^.

Taken together, DFINE provides a novel tool that allows for flexible inference of low-dimensional latent factors while capturing nonlinearities and enabling accurate neural, behavior, and manifold prediction. As such, DFINE can be used both for probing neural population activity across diverse domains of neuroscience and for developing future neurotechnology.

## Methods

### Neural data recordings

We performed our analyses on four diverse datasets containing distinct behavioral tasks, brain regions, and neural signal types to show the generalizability of the results.

#### Saccade task

A macaque monkey (Monkey A) performed saccadic eye movements during visually-guided oculomotor delayed response task^43^ (**Fig. 2a**). Each experimental session (13 sessions in total) consisted of several trials towards one of eight peripheral targets on a screen. All surgical and experimental procedures were performed in compliance with the National Institutes of Health Guide for Care and Use of Laboratory Animals and were approved by the New York University Institutional Animal Care and Use Committee. Trials began with the illumination of a central fixation square. The subject was trained to maintain its eyes on the square for about 500-800ms. After this baseline period, a visual cue was flashed for 300ms at one of the eight peripheral locations to indicate the target of the saccade (Target On event). After a delay, the central fixation square was extinguished, indicating the Go command to start the saccade (Go event). The subject was trained to perform the saccade (Saccade Start event) and maintain fixation on the target for an additional 300ms. A fluid reward was then delivered. The visual stimuli were controlled via custom LabVIEW (National Instruments) software. Eye position was tracked with an infrared optical eye tracking system (ISCAN) and from these positions some of the task events such as Saccade Start were identified. In this task, there are eight task conditions, each representing trials to one of the eight targets. During the task, LFP signals were recorded from lateral PFC with a semi-chronic 32-microelectrode array microdrive (SC32-1, Gray Matter Research). Raw LFP signals were low-pass filtered at 300 Hz and down-sampled to 20 Hz leading to a LFP observation every 50ms.

#### Naturalistic 3D reach-and-grasp task

A macaque monkey (Monkey J) performed a naturalistic reach-and-grasp task in a 50 × 50 × 50 cm^3^ workspace for a liquid reward across seven experimental sessions (**Fig. 2b**). All surgical and experimental procedures were performed in compliance with the National Institute of Health Guide for Care and Use of Laboratory Animals and were approved by the New York University Institutional Animal Care and Use Committee. During the task, the subject naturalistically reached for an object positioned on a wand, grasped it, released it and then returned its hand to a natural resting position^11^.

The wand was continuously moved around by the experimenter within a diverse spatial area in front of the subject^11^. The task was performed continuously in time without any instructions to isolate reach-and- grasp movement components. A total of 23 retroreflective markers were attached on the subject’s right arm and monitored using infrared and near-infrared motion capture cameras (Osprey Digital RealTime System, Motion Analysis Corp., USA) at a sampling rate of 100 frames *s*^−1^. We labeled 3D marker trajectories on the arm and hand (Cortex, Motion Analysis Corp., USA). The behavior variables were taken as the arm kinematics, i.e., the position and velocity of the wrist marker in the x, y and z directions. On each frame, motion capture camera data acquisition was synchronized to the neural recordings using a synchronization trigger pulse. The task lacked pre-defined trial structure or pre- defined target locations. Therefore, we identified the trial starts and ends from the velocity of the hand movement^11^, where the start of the trials was set to the start of the reach, and the end of the trials was set as the end of return and hold durations of the movement (**Fig. 2b**). To show condition-average visualizations, we partitioned the trials into two different conditions corresponding to left-ward or right- ward reaches along the horizontal axis in front of the subject, respectively. The horizontal axis was chosen for this division because it explained the largest variability in the reach locations.

Neural activity was recorded from M1, PMd, PMv and PFC in the left (contralateral) hemisphere with an array containing 137 microelectrodes (large-scale micro-drive system, Gray Matter Research, USA). Similar to our prior work^11^, we analyzed the pool of top 30 spiking channels sorted based on the individual channel behavior prediction accuracies. Spiking activity was calculated by counting the spikes in 10-ms bins and applying a Gaussian kernel smoother^50, 52, 73, 95, 96^ (with 30ms standard deviation), followed by down-sampling to have spiking activity observations every 50ms.

#### 2D random-target reaching task

A macaque monkey (Monkey T) performed a 2D random-target reaching task with an on-screen cursor in a total of three experimental sessions^45^ (**Fig. 2c**). All surgical and experimental procedures were consistent with the guide for the care and use of laboratory animals and approved by the institutional animal care and use committee of Northwestern University^45, 46^. The subject controlled the cursor using a two-link planar manipulandum while seated in a primate chair. Hand movements were constrained to a horizontal plane within a 20 × 20 cm^2^ workspace. The task consisted of several sections in each of which the subject performed 4 sequential reaches to random visual targets that appeared on the screen to receive a liquid reward (**Fig. 2c**). Within each section, after reaching the target and holding for a short period, the next target appeared in a pseudo-random location within a circular region (radius = 5-15 cm, angle = 360 degrees) centered on the current target. On average, the next target appeared approximately 200ms after the subject reached the current target. The task naturally consisted of non-stereotyped reaches to different locations on a 2D screen unlike traditional center-out cursor control tasks with stereotyped conditions^45^. Here, 2D cursor position and velocity in x and y directions were used as behavior variables. To show condition-average visualizations, we partitioned the reaches into 8 different conditions given the direction angle between the start and end point of the cursor trajectory (**Supplementary Fig. 7**). The angle of movement specifies the 8 conditions, which correspond to movement angle intervals of 0-45, 45-90, 90-135, 135-180, 180-225, 225-270, 270-315, and 315-360, respectively.

The subject was implanted with a 100-electrode array (Blackrock Microsystems) in PMd. After spike sorting, two sets of units were excluded from analysis in the original work^45^: 1) the units with low firing rates (smaller than 2 spike/s), 2) the units that had high correlations with other units. This led to 46, 49 and 57 number of units across different recording sessions. Spiking activity was calculated by counting the spikes in 10-ms bins and applying a Gaussian kernel smoother (with 30ms standard deviation), followed by down-sampling to have spiking activity observations every 50ms.

#### 2D grid reaching task

A macaque monkey (Monkey I) performed a 2D grid reaching task by controlling a cursor on a 2D surface in a virtual reality environment^47, 48^ (**Fig. 2d)**. All animal procedures were performed in accordance with the U.S. National Research Council’s Guide for the Care and Use of Laboratory Animals and were approved by the UCSF Institutional Animal Care and Use Committee^47, 48^. Circular targets with 5mm visual radius within an 8-by-8 square grid or an 8-by-17 rectangular grid were presented to the subject and the cursor was controlled with the subject’s fingertip position. Fingertip position was monitored by a six-axis electromagnetic position sensor (Polhemus Liberty, Colchester, VT) at 250 Hz and then non-causally low-pass filtered to reject sensor noise (4th order Butterworth filter with 10 Hz cutoff). The subject was trained to acquire the target by holding the cursor in the respective target-acceptance zone, a square of 7.5mm edge length centered around each target, for 450ms. After acquiring the target, a new target was drawn from the possible set of targets. In most sessions, this set was generated by replacement, i.e., the last acquired target could be drawn as the new target. However, the last acquired target was removed from the set in some sessions. Even though there was no inter-trial interval between consecutive reaches, there existed a 200ms “lockout interval” after target acquisition where no new target could be acquired. 2D cursor position and velocity in x and y directions were used as behavior variables. To show condition-average visualizations, we partitioned the reaches into 8 different conditions based on their direction angle as in the 2D random-target reaching task.

One 96-channel silicon microelectrode array (Blackrock Microsystems, Salt Lake City, UT) was implanted into the subject’s right hemisphere (contralateral) M1. Total of seven sessions (sessions 20160622/01 to 20160921/01) were used in our manuscript^39^. We analyzed the pool of top 30 neurons sorted based on the individual neuron behavior prediction accuracies. Spiking activity was calculated by counting the spikes in 10-ms bins and applying a Gaussian kernel smoother (with 30ms standard deviation), followed by down-sampling to have spiking activity observations every 50ms.

### DFINE model

We develop a neural network architecture that allows for accurate nonlinear description similar to deep neural networks, but in a manner that also enables flexible inference similar to LDM. To do so, we develop a new neural network in which we introduce two distinct sets of latent factors for *n*_y_- dimensional neural population activity **y**_*t*_ ∈ ℝ^*n*y×1^: dynamic latent factors **x**_*t*_ ∈ ℝ^*n*x×1^ and manifold latent factors **a**_*t*_ ∈ ℝ^*n*a×1^, where *n*_x_ and *n*_a_ are the factor dimensions/hyperparameters to be picked. The key idea here is to incorporate a middle noisy manifold layer **a** between the dynamic latent factors **x** and neural observations ***y***, which allows us to separate the model into a nonlinear manifold component and a linear dynamical component evolving on this nonlinear manifold (**Fig. 1a**). This separation plays a key role in enabling the new flexible inference properties of the network to optimally perform: 1) recursive causal inference (filtering), 2) non-causal inference with all observations (smoothing), and 3) inference in the presence of missing observations. Specifically, the separation enables flexible inference using a Kalman filter for the bottom dynamic part of the model in **Fig. 1a** that infers the dynamic factors **x** from the manifold factors **a** (**Fig. 1b**). We now describe the network architecture consisting of the dynamic and manifold factors.

First, the dynamic latent factor evolves in time with a linear Gaussian model:

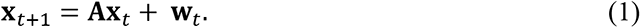

where **W**_*t*_ ∈ ℝ^*n*_x_×1^ is a zero-mean Gaussian noise with covariance matrix **W** ∈ ℝ^*n*_x_×*n_x_*^ and **A** ∈ ℝ^*n*_x_×*n_x_*^ is the state transition matrix. The manifold latent factor **a**_*t*_ is related to the dynamic latent factor **x**_*t*_ as:

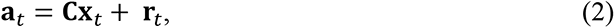

where ***C*** ∈ ℝ^*n_a_*×*n_x_*^ is the emission matrix and ***r***_*t*_ ∈ ℝ^*n_a_*×1^ is a white Gaussian noise with covariance matrix ***R*** ∈ ℝ^*n_a_*×*n_a_*^. Equations (1) and (2) together form an LDM with **a**_*t*_ being the Gaussian noisy observations. We denote the parameter set of this LDM by *Ψ* = {**A**, **W**, **C**, **R**, μ_0_, **Λ**_0_}, where **μ**_0_ and **Λ**_0_are the initial estimate and covariance of the dynamic latent factors, respectively.

Second, to model nonlinearities, we use autoencoders to learn the mapping between neural observations **y**_*t*_ and manifold latent factors **a**_*t*_. In general, autoencoders are static generative models made of two parts: the encoder that maps the observations to a bottleneck representation and the decoder that takes this bottleneck representation to the observations. Here, autoencoder observations are the neural observations and autoencoder bottleneck representation is given by the manifold latent factors.

We model the decoder part as a nonlinear mapping f_*θ*_(⋅) from manifold latent factors to neural observations:

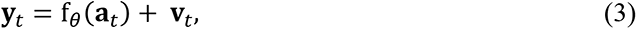

where *θ* are parameters and ***v***_*t*_ ∈ ℝ^*n*y×1^ is a white Gaussian noise with covariance ***V*** ∈ ℝ^*n*y×*n*y^. We model nonlinear mappings with MLPs as they are universal approximators of any nonlinear function under mild conditions^40^. Equations (1)-(3) together form the generative model (**Fig. 1a**). For inference (next section), we also need the mapping from **y**_*t*_ to **a**_*t*_, which we characterize as:

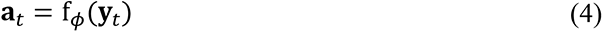

where f_*ϕ*_(⋅) represents the encoder in the autoencoder structure and is parameterized by another MLP (**Fig.** 1**a**).

#### The inference problem

Using the model in equation (1)-(4), we need to infer both the manifold and dynamic latent factors from neural observations **y**_1:*T*_, where *T* is the total number of time steps for the observations. We use the subscript *t*|*k* to denote the inferred latent factors at time *t* given **y**_1:*k*_. Therefore, *t*|*t* denotes filtering (causal) inference given **y**_1:*t*_ and *t*|*T* denotes smoothing (non-causal) inference given **y**_1:*T*_. As an intermediate step called nonlinear manifold embedding, we first directly but statically get an initial estimate of **a**_*t*_ based on **y**_*t*_ from equation (4) as **â**_*t*_ = f_*ϕ*_(**y**_*t*_) to provide the noisy observations of the dynamical model, i.e. **â**_*t*_, in **Fig. 1b**. Having obtained **â**_*t*_, we can now use the dynamical part of the model in equations (1) and (2) to infer **x**_*t*|*t*_ with Kalman filtering from **â**_1:*t*_, and infer **x**_*t*|*T*_ with Kalman smoothing^97^ from **â**_1:*T*_. We can then infer the manifold latent factor as **a**_*t*|*t*_ = ***C*****x**_*t*|*t*_ and **a**_*t*|*T*_ = ***C*****x**_*t*|*T*_based on equation (2). Note that **a**_*t*|*t*_ is inferred not only based on the neural observations but also based on the learned dynamical model using the Kalman filter, and thus this inference aggregates information over time dynamically.

The inference above has the following major advantages compared with prior generative neural network models that train a non-causal inference network and use all observations in a trial. First, we can tractably infer latent factors recursively in real-time, i.e., use the current inferred dynamic latent factors **x**_*t*|*t*_ to calculate the next step’s inferred value **x**_*t*+1|*t*+1_. Second, we can handle missing observations by doing only forward prediction with the Kalman predictor at time-steps when observations are missing, without any need to impute 0 value for these missing observations as done previously^11, 28, 98^. Third, we can perform both causal filtering and non-causal smoothing inference with the same learned model.

#### The learning problem

Neural network model parameters are learned to optimize a cost function. When stochastic noise variables exist^4, 30, 64^, this cost is typically taken as the ELBO. But optimizing ELBO can be difficult^68–71^ and the direct goal of dynamical modeling is to predict neural and/or behavior dynamics. Thus, we instead define our cost as the k-step-ahead prediction error in predicting neural observations *k* time- steps into the future, i.e., the error between ***y***_*t*+*k*_ and its prediction from ***y***_1:*t*_ denoted by ***y***_*t*+*k*|*t*_. What allows us to use this cost is that our model enables efficient recursive inference/prediction to compute the k-step-ahead prediction error (**Fig. 1b**) in the presence of noise; this is because we can run all forward predictions with a single run of Kalman filtering. Thus, our cost *L* is a function of all parameters as:

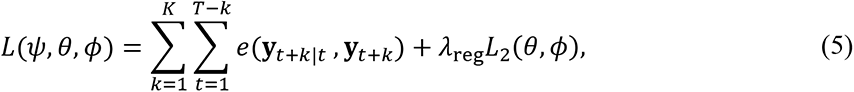

where *K* denotes the maximum horizon for future-step-ahead prediction and *e*(⋅,⋅) denotes the error measure. *T* is the length of the time-series, i.e., the length of each batch in the mini-batch gradient descent^99^. *L*_2_(*θ*, *ϕ*) is an *L*_2_ penalty for the autoencoder parameters {*θ*, *ϕ*} to prevent overfitting with regularization hyperparameter *λ*_reg_^99^. In practice, we use root mean-squared error (RMSE) for the error measure *e*(⋅,⋅).

#### Supervised DFINE learning

In the unsupervised DFINE, latent factors are optimized to be predictive of neural population activity. To address situations in which neural dynamics of a specific behavior are of interest, we develop a new learning algorithm for DFINE that aims to extract latent factors that are more predictive of that behavior. We call the resulting algorithm supervised DFINE. In particular, when continuous behavior variables ***z***_*t*_ ∈ ℝ^*n*z×1^ are available during training, we use them to supervise the training and learn latent factors that are predictive of not only neural observations but also behavior variables. This is done by adding an auxiliary neural network, termed mapper MLP network, to the DFINE graphical model during training only (**Fig. 1a**); this mapper network maps the manifold latent factor **a**_t_ to behavior variables ***z***_*t*_ during training (the link from **a** to ***z*** in **Fig. 1a**) and is written as ***z***_*t*_ = *f*_*γ*_(**a**_*t*_) + ***q***_*t*_, where ***q***_*t*_ ∈ ℝ^*n*z×1^ is white Gaussian noise. Now to motivate the network to learn latent factors that are predictive of both behavior and neural observations, we add the behavior prediction error to the cost function in equation (5) as follows:

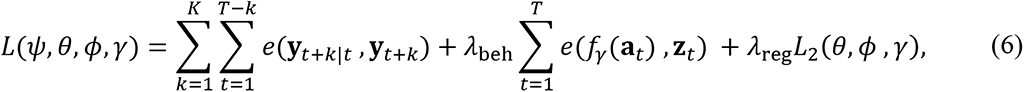

where *λ*_beh_ is a hyperparameter. Larger values of *λ*_beh_ put more emphasis on behavior prediction vs. neural prediction and vice versa. Note that the parameters of the auxiliary MLP (*γ*) are also added to the regularization term in the cost function.

We emphasize that after training of supervised DFINE is completed, the mapper MLP is discarded and not used for inference. The inference of latent factors remains identical to that in unsupervised DFINE, and is done purely based on neural observations and independent of behavior variables (**Fig. 1b**). The only difference is that supervised DFINE’s learned model has different parameter values given that its learning algorithm is distinct as described above.

#### DFINE learning: hyperparameters and details

Given a set of observation, we learn the model parameters by minimizing the cost function in (5). We used MLP architectures containing 3 hidden layers each with 32 units for *f*_*θ*_(⋅) and *f*_*ϕ*_(⋅) in decoder and encoder parts of the model, respectively. The activation function used for the units was set to tanh(⋅). We used back propagation with mini-batch gradient descent implemented by ADAM optimizer^100^ to learn the model parameters and we continued the training for 300 epochs. We used 0.02 for the initial learning rate of the ADAM optimizer. We set the maximum horizon for future-step-ahead prediction *K* such that the future predictions cover at least 100ms into the future, therefore we set *K* = 2 − 4 to optimize the cost function across various datasets (note our time step is 50ms). We use the regularization parameter *λ*_reg_ = 10 to prevent overfitting. *λ*_beh_ was set to 100 across all the supervised DFINE models to put emphasis on the improved behavior prediction accuracy.

### Evaluation using five-fold cross-validation

For all analyses in this work, we performed five-fold cross-validation, where we divide the data into five equal-sized folds, use 4 folds as the training set to learn the model, and leave one fold out for the test set to evaluate the learned models. Below, we expand on the evaluation metrics used in this manuscript.

#### Behavior prediction accuracy

##### Saccade task

For any method, we quantified the behavior prediction accuracy of the latent factor time-series during movement periods by calculating the target classification accuracy using these factors (**Fig. 2a**). To address the inter-trial variability in the length of movement periods so that the classifier can be applied to latent factors in any trial, we performed an identical preprocessing step for the latent factors from all methods. For this preprocessing, we linearly interpolated the latent factor time-series duration to 400 datapoints and uniformly sampled 10 datapoints (i.e., every 40 datapoints after interpolation). After this preprocessing and by flattening the processed factors, we obtained the classification features with dimension equal to 10 × latent factor dimension. For training and test trials, we used these processed features to learn classifiers and perform classification. In addition, to compute the performance given the limited number of trials in the training and test sets, for all methods, we averaged the classification accuracies of all binary classifiers (for each class vs. another class) rather than performing an 8-class classification. We used nonlinear support vector machines (SVM) with Gaussian kernels to perform the binary classification. The width or standard deviation of the Gaussian kernel for each training fold was picked based on inner cross-validation. To assess the target classification accuracy, we used AUC of the receiver’s operating curve^41^ for all binary classifiers and computed the mean performance for each test cross-validation fold.

##### Motor datasets

Here behavior variables were continuous. For any method, to quantify the behavior prediction accuracy, we learned an MLP regression model from the smoothed latent factors to the observed behavior variables in the training set. In the test set, we used the learned MLP regression model to get the predicted behavior variables from the latent factors. We used Pearson’s correlation coefficient (CC) to quantify the behavior prediction accuracy in each test cross-validation fold.

#### Neural prediction accuracy

We quantify the neural prediction accuracy by calculating how accurately models predict neural observations one-step-ahead into the future from their own past. For the DFINE model, we use **y**_*t*+1|*t*_ = f_*θ̂*_**a**_*t*+1|*t̂*_ as the one-step-ahead neural prediction. For LDM, neural prediction is given by the classic Kalman predictor^89, 98^. However, sequential autoencoders (SAE) do not perform one-step-ahead prediction during the trials as they have a non-causal inference architecture and need to observe all neural observations to infer the latent factors and reconstruct the neural observations (see **Discussion**). We thus give SAEs an advantage by allowing them to use all neural observations (instead of just past observations) and thus to perform neural reconstruction instead of one-step-ahead neural prediction. We also compared DFINE’s neural reconstruction accuracy given by **y**_*t*|*T*_ = f_*θ̂*_**a**_*t*|*T̂*_ to SAE’s neural reconstruction accuracy in **Supplementary Fig. 2** with similar conclusions. We use Pearson’s correlation coefficient (CC) to quantify the neural prediction accuracy in each test cross-validation fold.

#### Topological data analysis (TDA) metrics

To quantify how robustly models identify the latent manifold structure in single-trials, we applied TDA^42^ on smoothed latent factors. TDA uses persistent homology^42^ to find multi-dimensional holes (e.g. 1D hole is a ring, 2D hole is a 2D void) in data by growing the radius of ***ϵ***-balls around datapoints, which are connected when they touch. TDA finds the manifold type by counting the number of persistent multi-dimensional holes in the data-manifold. For example, a torus has one 2D hole and two 1D holes^42^. We run TDA on smoothed single-trial latent factors of the learned models in the cross- validation test set and assess the most persistent hole’s birth and length. The most persistent hole is the hole that lasts the longest. The birth happens at the ***ϵ*** value at which the hole appears (smaller values correspond to earlier births) and the length is the ***ϵ*** interval for which the hole lasts. To assess robustness in single-trials, we ask for which model TDA finds holes that are born earlier and last longer, i.e., are more persistent: the sooner a hole is born (at shorter radius) and the longer it lasts (at longer radius), the more prevalent/robust it is in the latent factors extracted. To take into account scaling differences in the latent space of each method, we z-score single-trial latent factors in each dimension before running TDA. To visualize the TDA results, for the most persistent holes, we plot their lengths vs. births. On this plot, the optimal frontier is the top left area of the plot indicating earlier births and longer lengths. To aggregate the results in each test cross-validation fold, we average the birth and length values of TDA for single-trial latent factors in that test fold.

### Implementation details for benchmark methods

**SAE:** For SAE, we use the architecture named LFADS^4^, which is a common benchmark nonlinear model of neural population activity^12, 29, 32^. LFADS is a RNN-based variational SAE^4^. LFADS takes fixed-length segments of data as input and encodes each segment into a bottleneck latent factor, which serves as the initial condition for the decoder’s network (i.e., generator). Given this initial condition, the generator RNN propagates the latent states, generates a factor time-series, which then reconstructs a smoothed copy of the input data segment. We train LFADS using the publicly available source code^4^ and using hyperparameters in row 2 of the Supplementary table 1 in the original manuscript^4^ because the number of trials in the dataset associated with row 2 was closest to our datasets. The dimensionality of the generator RNN’s latent state in LFADS represents its dynamical memory and the number of values used at any given time-step to generate the dynamics/states of the next time-step, thus representing its dynamics state dimension^12^. We keep the generator RNN’s latent state dimension (and thus the initial condition latent factor dimension) high enough and set it to 64. This gives an advantage to LFADS in terms of modeling the dynamics because it gives LFADS a higher dynamics state dimension of 64 compared to DFINE that has this dimension at 16. We use a Gaussian observation distribution to train LFADS so that, like DFINE, it can be applied for all neural modalities considered here whether spike counts or LFP. We use the LFADS factor time-series as the latent factors of LFADS in our analyses/comparisons as was done in the original manuscript^4^.

LFADS can only be trained on 3D data tensors (trial × time × observation dimension) given its SAE structure. Thus, for its model training and inference, we split the continuous data into smaller 1s segments to create 3D data tensors in both training and test sets^12, 84^. We then trained the LFADS model with the training set and performed inference in each 1s segment of data. We finally got the smoothed latent factors for the full duration of training and test sets by concatenating the inferred latent factors across segments. Similar to the original paper^4^, we ran the inference 50 times – as LFADS inference is stochastic given its variational autoencoder format^4^ – and averaged the inferred latent factors from these 50 realizations.

**LDM**: Similar to our prior work^12^, we train LDM models using a publicly available Python package that performs subspace identification^12^. Throughout the manuscript, we use the states of the LDM model as the latent factors. The latent factors are inferred using Kalman filtering and smoothing.

### Numerical simulations

We validate the DFINE model on numerical simulations first. Since prior studies on neural population activity have shown significant evidence for rotatory dynamics and ring-like/Toroidal manifolds^6, 10, 15, 49^, we simulate the nonlinearity using 3 different manifold types to show generality: ring- like, Torus and Swiss roll manifolds. We simulate nonlinear trajectories over these manifolds by first generating a driven-walk using a linear dynamical model on the manifold local coordinates and then embedding these coordinates in 3D cartesian space using the manifold equations (see **Supplementary Note 1**). To get the neural observations, we generate 40D output signals by first applying a random output emission matrix (of size 40 × 3) on the 3D trajectories and then adding additive white Gaussian noise to the 40D signal to realize noisy observations. Without loss of generality and only for illustration purposes, we keep the first 3 dimensions of the observations the same as the 3D trajectories (i.e., use identity output submatrix for the first 3 dimensions) so that we can illustrate the first 3 dimensions for our 3D visualizations without any linear distortions. We emphasize that all quantifications are calculated with 40D observations (see **Supplementary Note 1** for more details). We simulate 30 different sessions (10 from each manifold type), where we randomly pick the white Gaussian noise standard deviations. In each session, we simulate 250 trials, each containing 200 time-steps. Our validations are all based on five-fold cross-validation on these trials.

In addition to visualization, we quantify the success of learning with the convergence of one-step-ahead neural prediction error in the learned models to that of the true model. We calculate the one-step-ahead prediction error using normalized root mean-squared error (NRMSE); this normalization allows for pooling the results across sessions given potential scaling differences across various simulated sessions. Given a one-dimensional signal *y*_*t*_ and its prediction *ŷ*_*t*_, NRMSE is calculated as:

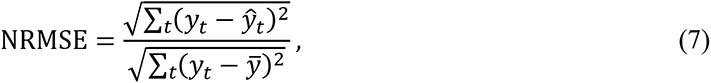

where *ŷ* is the mean of the signal. We calculate the NRMSE for each dimension and report the mean across dimensions for the 40D observations in our case. To calculate the one-step-ahead prediction error, we calculate the NRMSE between the true observations (***y***_*t*_) and their predictions one-step into the future (***ŷ***_*t*|*t*−1_). For the true models, we calculate the one-step-ahead predictions using Unscented Kalman filtering (UKF)^101^ as the true manifolds have nonlinearities that we can write analytically for the true model and write a UKF for.

### Inference analyses with missing observations

We assess/visualize the inference in the presence of missing observations across various observed datapoint ratios. Observed datapoint ratio, denoted by *ρ*, quantifies the ratio between the number of observed datapoints versus the total number of datapoints. We train both the DFINE and SAE models on fully observed training sets, and then test their inference on test sets with missing observations. For simulation analyses, we vary *ρ* from 0.05 to 1 by randomly dropping datapoints in each test trial, infer the latent factors in the presence of missing/dropped observations, and then quantify the error between the true and reconstructed manifold trajectories via filtering/smoothing. For motor datasets, we introduced missing observations by randomly and uniformly dropping neural observations with various observed datapoint ratios (*ρ* ranging from 0.2 to 1). Since the motor datasets contained continuous recordings and to make sure that we dropped datapoints uniformly throughout the duration of time- series, we randomly dropped (1 − *ρ*) × 100 datapoints in every 100 time-steps of the neural observation time-series.

Using the learned DFINE models in the test set, we inferred the latent factors at all time-steps even though observations were missing at some random time-steps. From these latent factors, the behavior variables were predicted in the test set using the learned MLP models and thus the cross-validated behavior prediction accuracy was computed. Note that, even though the observations are missing at random time-steps, the latent factors and thus behavior variables are inferred at all time-steps. In a control analysis, we also perform inference with SAE in the presence of missing observations as described above. For SAEs, we impute the missing observations with zeros^33, 34^ since SAE’s generator RNN is designed to take inputs at every time-step. Given the SAE models that are trained on fully observed training sets, we infer the latent factors in the test sets with missing observations, predict behavior variables with learned MLP models and compute the cross-validated behavior prediction accuracy.

### Statistical analysis

For all analyses in this work, significance was declared if *P* < 0.05. All statistical tests were performed with non-parametric Wilcoxon signed-rank tests.

## Acknowledgements

The authors acknowledge support of NIH Director’s New Innovator Award DP2-MH126378 (to M.M.S.), NIH R01MH123770 (to M.M.S.), and NSF CRCNS Award IIS 2113271 (to M.M.S. and B.P.).

## Author Contributions

H.A. and M.M.S. conceived the study and developed the new algorithms. H.A. performed all the analyses except for the grid task. E.E. performed the analysis for the grid task. H.A. and M.M.S. wrote the manuscript. B.P. designed and performed the experiments for two of the nonhuman primate datasets.

M.M.S. supervised the work.

## Supplementary Figures

**Supplementary Figure 1.**
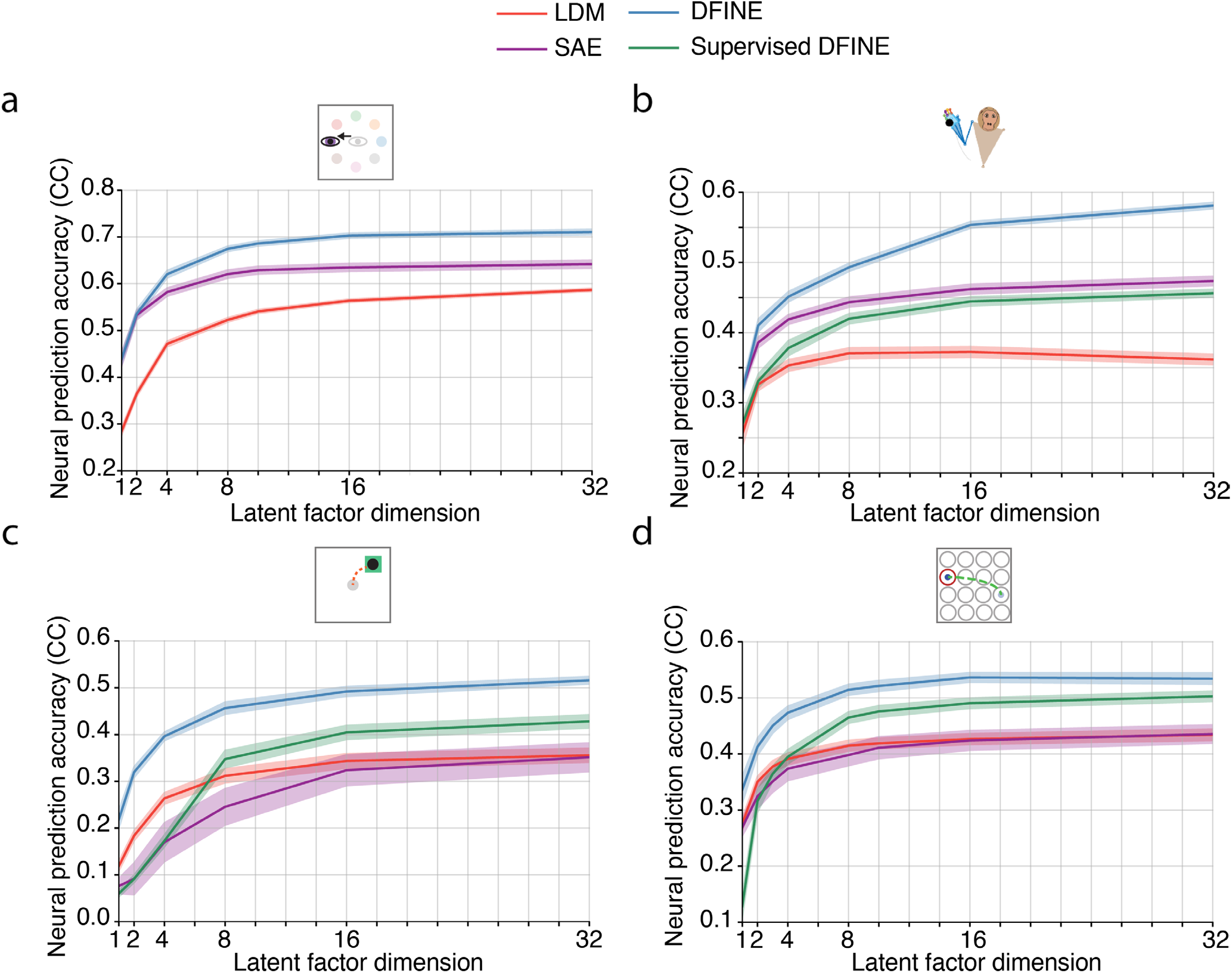
Neural prediction accuracy as a function of the latent factor dimension is shown for all datasets and methods. Solid lines show the mean neural prediction accuracy across sessions and cross validation folds for the (a) saccade task (observation dimension *n*_y_ = 32), (b) 3D naturalistic reach-and-grasp task (observation dimension *n*_y_ = 30), (c) 2D random-target reaching task (observation dimension *n*_y_ = 46 − 57), and (d) 2D grid reaching task (observation dimension *n*_y_ = 30). The shaded areas represent the 95% confidence bound. For SAE, the dimension shown is the factor dimension and the dynamic dimension (generator RNN’s latent state dimension and initial condition dimension) is always 64.

**Supplementary Figure 2.**
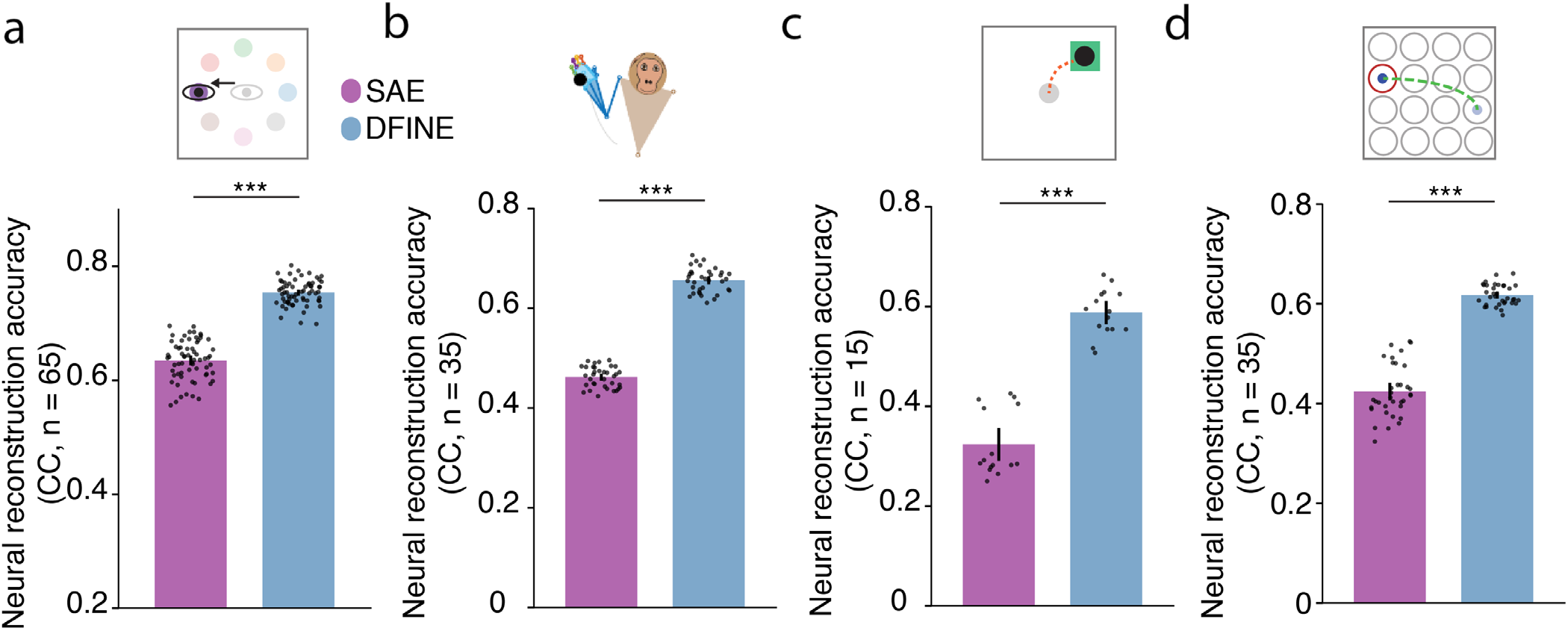
DFINE’s neural reconstruction accuracy with smoothing is also better than that of SAE. Figure convention is as in Fig. 4. The neural reconstruction accuracy with smoothing is shown for the (a) saccade task, (b) 3D naturalistic reach-and-grasp task, (c) 2D random-target reaching task, and (d) 2D grid reaching task.

**Supplementary Figure 3.**
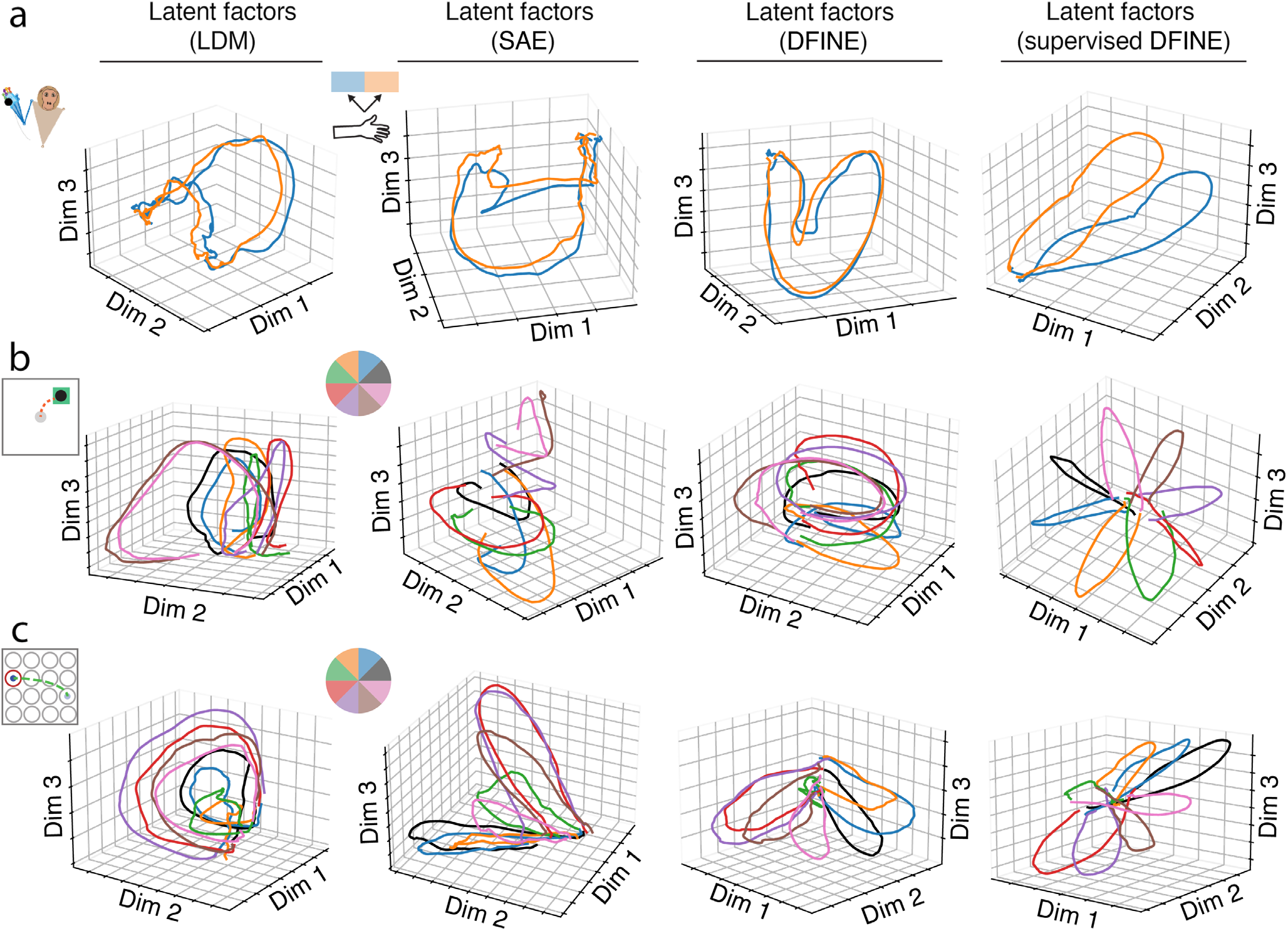
Example latent factor trajectories for the motor datasets. Figure convention for the conditions is as in Fig. 6. The condition-average latent factor trajectories are shown for all methods in the (a) 3D naturalistic reach-and-grasp task, (b) 2D random-target reaching task, and (c) 2D grid reaching task. We observed a ring-like manifold structure during movement periods and DFINE more robustly identified this ring-like structure in single-trials as evidenced by the TDA results in Figs. 4 and 5.

**Supplementary Figure 4.**
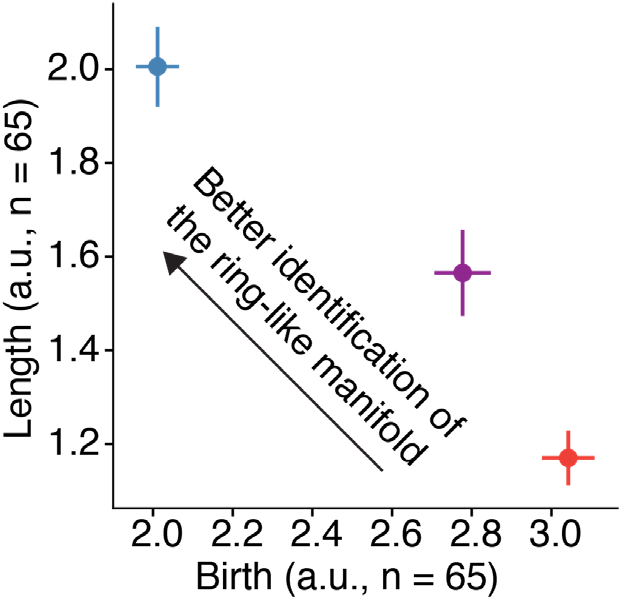
DFINE more robustly extracts the ring-like manifold structure in single-trials during the preparation period of the saccade task. TDA analysis on single-trial latent factors during the preparation period is shown. TDA’s most persistent 1D hole had a significantly earlier birth and lasted significantly longer for DFINE compared to LDM and SAE (*P* < 5 × 10^−4^, one-sided Wilcoxon signed-rank test, n = 65). Figure convention is the same as Fig. 4e. Example condition-average and single-trial latent factor trajectories during the preparation period are shown in Fig. 4a.

**Supplementary Figure 5.**
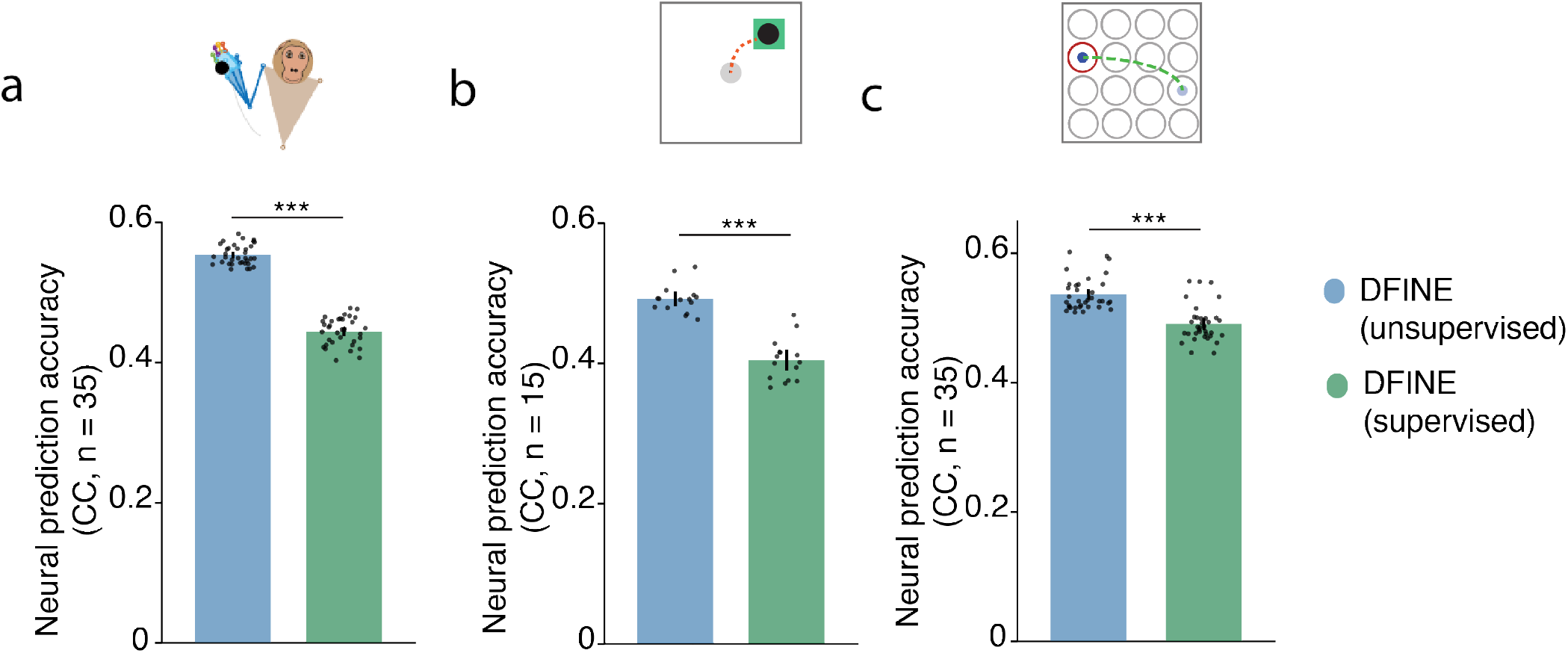
Supervised DFINE had lower neural prediction accuracy compared to unsupervised DFINE as expected as the former optimizes for both neural and behavior prediction rather than just for neural prediction. Figure convention is as in Fig. 6b. Neural prediction accuracies are shown for the: (a) 3D naturalistic reach-and-grasp task, (b) 2D random-target reaching task, and (c) 2D grid reaching task.

**Supplementary Figure 6.**
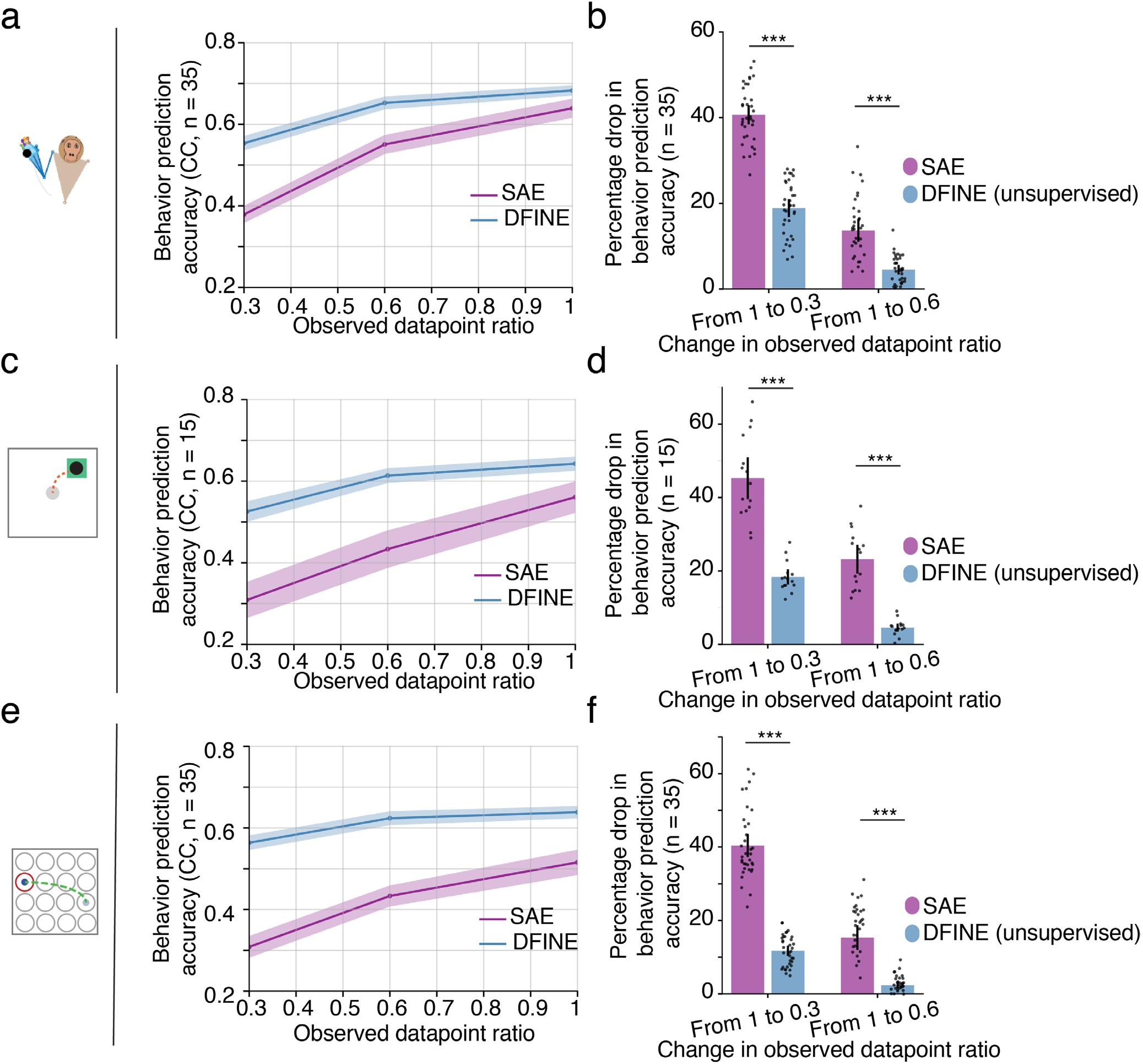
DFINE outperforms SAE in the presence of missing observations and this improvement grows with more missing samples. (a) DFINE and SAE’s behavior prediction accuracy across various observed datapoint ratios in the 3D naturalistic reach-and-grasp task. Figure convention is similar to that in Fig. 7. Given models trained on fully observed neural observations, we inferred latent factors in the test set that had missing observations. SAE did so by imputing missing observations in the test set to zero as done previously^33, 34^, whereas DFINE did so through its new flexible inference method. We then used the inferred factors in the test set to predict behavior variables. This process was done at 0.3 and 0.6 observed datapoint ratios. For both models, we show the behavior prediction accuracy of the smoothed latent factors. (b) The percentage drop in the behavior prediction accuracy of DFINE and SAE as we vary the observed datapoint ratio from 1 to 0.3 and from 1 to 0.6. The percentage drop in behavior prediction accuracy of DFINE is significantly lower than that of SAE, showing that DFINE can better compensate for missing observations. Figure convention for bars, dots and asterisks is similar to that in Fig. 4. Similar results held for the 2D random target reaching task (c,d), and for the 2D grid reaching task (e,f).

**Supplementary Figure 7.**
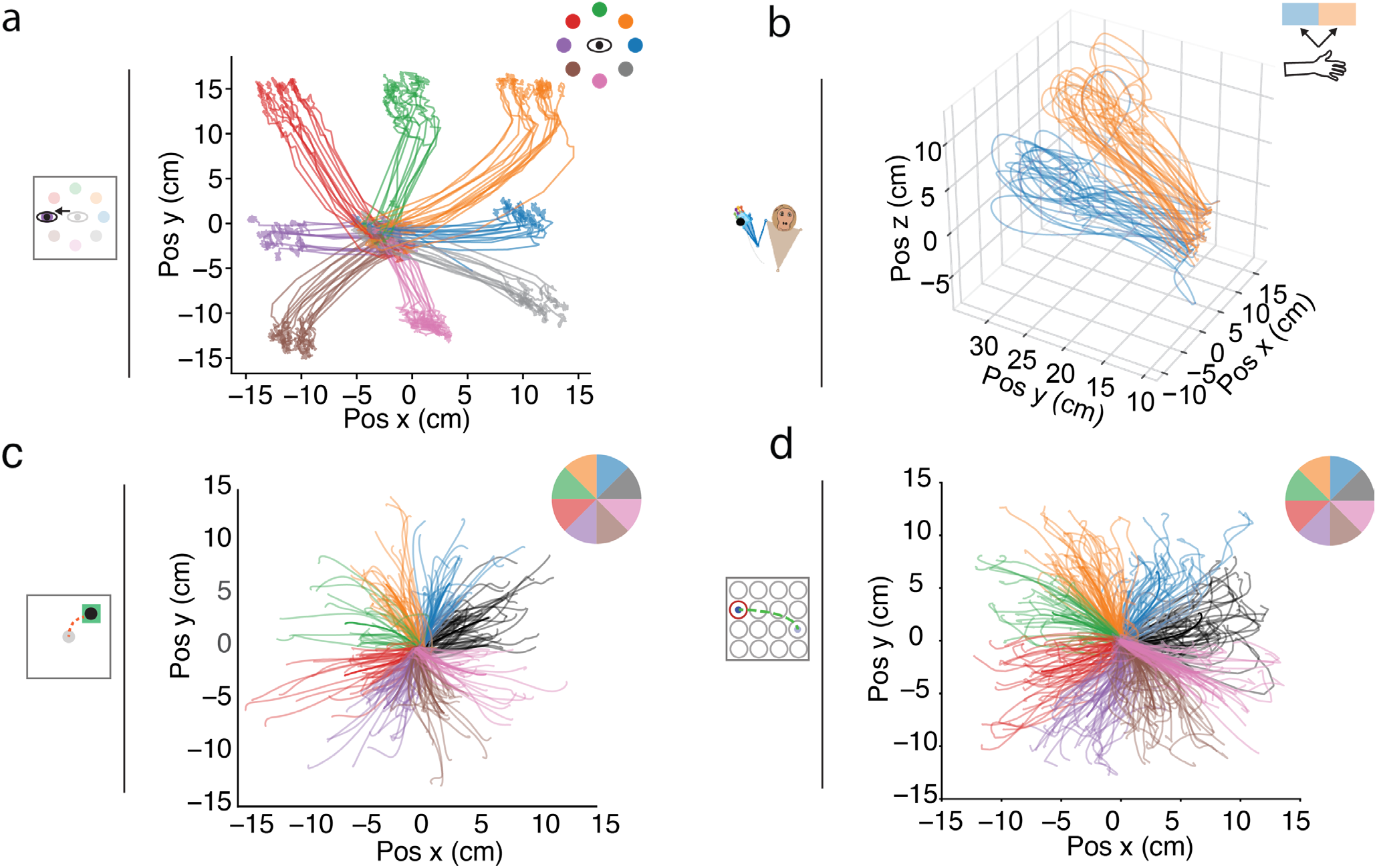
Example behavior trajectories for the four experimental datasets. (a) Eye movement trajectories for the saccade task. Each color represents one target, i.e., condition. (b) 3D hand movement trajectories for the 3D naturalistic reach-and grasp task. Each color represents one condition, i.e., movement to left or right. 2D cursor trajectories for the (c) 2D random-target reaching task and (d) 2D grid reaching task are shown, when shifted in space to start from the center. Each condition is shown with a different color and represents reaches that have similar direction angles. Regardless of start or end position, the angle of movement specifies the 8 conditions, which correspond to movement angle intervals of 0-45, 45-90, 90-135, 135-180, 180-225, 225-270, 270-315, and 315-360, respectively.

## Supplementary Notes

### Supplementary Note 1. Details of numerical simulations

We simulate 3 different manifold types including ring-like, Torus and Swiss roll manifolds. Here, we expand on the numerical simulations.

#### Manifold equations

For each manifold type, we first get the 3-dimensional (3D) manifold embeddings, which specify how the manifold is embedded in 3D Cartesian space and are used as visualizations (**Supplementary Note Fig. 1**). We then transform these embeddings using a random output emission matrix to a 40D space to get the neural observations (see next section). To get the 3D manifold embeddings, each manifold type has its own equation, which we expand on in the following sections. Below, we denote the 3 dimensions of the 3D embeddings as e_1_, e_2_ and e_3_.

**Ring-like manifold:** We generate the ring-like manifold embeddings from the 1D ring manifold coordinate (d_θ_), which is an angle between 0-2*π*, using the following equations (**Supplementary Note Fig. 1a**):

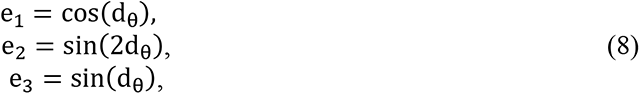

**Torus manifold:** We generate the Torus manifold embeddings from the 2D Torus manifold coordinates d_r_ and d_R_ (**Supplementary Note Fig. 1b**). d_r_ is the coordinate – angle between 0-2*π* – for the minor circle which is the inner circle representing the Torus’s tube. d_R_ is the coordinate – angle between 0-2*π* – for the major circle which is the outer circle on which the Torus’ tube evolves. R and r are the radius values for the minor and major circles. We get the 3D manifold embeddings as:

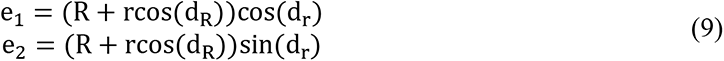

Without loss of generality, we use R = 4 and r = 1.5 for the major and minor radii, respectively.

**Swiss roll manifold:** The below equation generates the 3D Swiss roll manifold embeddings from the 2D Swiss-roll coordinates d*_r_* and d_h_, which are its circular and height coordinates, respectively (**Supplementary Note Fig. 1c**):

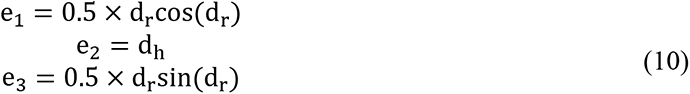

**Supplementary Note Figure 1.**
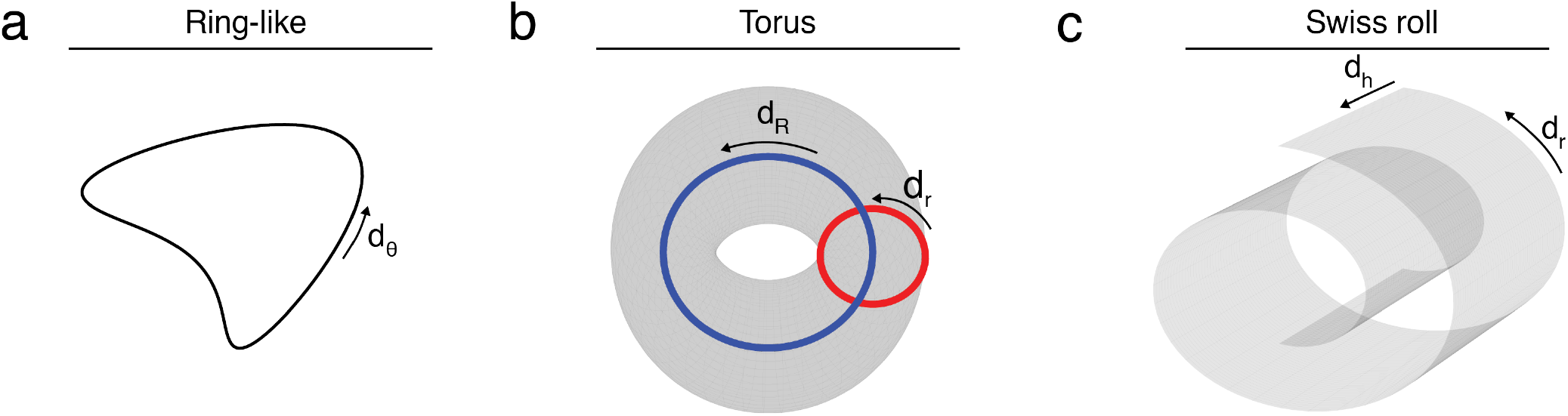
Visualizations of the manifolds. 3D manifold embeddings are shown for **(a)** ring-like, **(b)** Torus, and **(c)** Swiss roll manifolds.

#### Generating neural trajectories over manifolds

We denote the vector manifold coordinates of the trajectories at each time step by **d***_t_*. Thus **d***_t_* is [d_θ_], [d_r_; d_R_] and [d_r_; d_h_] for ring, Torus and Swiss roll manifolds, respectively. We generate trajectories over the manifolds by first generating a walk on the manifold’s local coordinate space with a linear dynamical equation:

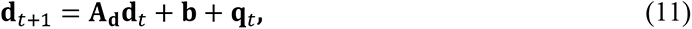

where **A_d_** is the diagonal state transition matrix with eigenvalues of 0.99, **b** is the input term to drive the trajectory at each time-step (set as 0.2), and ***q***_*t*_ is the white Gaussian noise with covariance **Q**. The standard deviation of **q**_*t*_ is randomly chosen between [0.01, 0.1]. After generating the trajectories from equation (11), we embed the manifold coordinate vector **d**_*t*_ within 3D Cartesian space to get the embeddings **e**_*t*_ with equations (8)-(10) and we finally get the neural observations from:

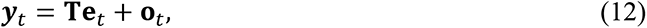

where **T** is the output emission matrix and **o**_*t*_ is a white Gaussian noise with covariance matrix O. The matrix **T** ∈ ℝ^40×3^ has its first 3 rows chosen as [1,0,..,0], [0,1,0,…,0], [0,0,1, 0,…,0] to form an identity transformation for the first 3 dimensions of the manifold (for visualization purposes only), and the rest of the 37 rows are randomly chosen with elements between [−5,5]. The standard deviation of **o**_*t*_ is randomly chosen between [5,25].

## Notes

### Competing Interest Statement

The authors have declared no competing interest.

